# NANOS2 suppresses the cell cycle by repressing mTORC1 activators in embryonic male germ cells

**DOI:** 10.1101/2020.09.23.310912

**Authors:** Ryuki Shimada, Hiroko Koike, Takamasa Hirano, Yumiko Saga

## Abstract

During murine germ cell development, male germ cells enter the mitotically arrested G0 stage, which is an initial step of sexually dimorphic differentiation. The male specific RNA-binding protein NANOS2 has a key role in suppressing the cell cycle in germ cells. However, the detailed mechanism of how NANOS2 regulates the cell cycle remains unclear. Using single-cell RNA sequencing (scRNA-seq), we extracted the cell cycle state of each germ cell in wild-type and *Nanos2*-KO testes, and revealed that *Nanos2* expression starts in mitotic cells and induces mitotic arrest. We also found that NANOS2 and p38 MAPK work in parallel to regulate the cell cycle, suggesting that several different cascades are involved in the induction of cell cycle arrest. Furthermore, we identified *Rheb*, a regulator of mTORC1, and *Ptma* as possible targets of NANOS2. We propose that the repression of the cell cycle is a primary function of NANOS2 and that it is mediated via the suppression of mTORC1 activity by repressing *Rheb* in a post-transcriptional manner.

## Introduction

The sexually differentiated gametes sperm and eggs originally differentiated from common precursors, primordial germ cells (PGCs). PGCs are specified at the posterior end of gastrulating embryos at E7.25 and migrate towards the gonads in mice (Hayashi et al. 2007; Anderson et al. 2000). After colonizing the gonads, PGCs start sex-specific differentiation by responding to factors supplied from surrounding somatic cells. In ovaries, retinoic acid (RA) from the mesonephros induces the expression of meiosis initiators, *Stimulated by RA gene 8* (*Stra8*) and *Meiosis initiator* (*Meiosin*), in germ cells to start meiosis (Koubova et al. 2006; Anderson et al. 2008; Ishiguro et al. 2020). On the other hand, in testes, RA is degraded by CYP26B1 produced in somatic cells (Saba et al. 2014b; Bowles et al. 2006), thus germ cells cannot enter meiosis. Instead, male germ cells enter mitotic arrest from E13.5 and most are arrested at the G1/G0 phase at E15.5 (Western et al. 2008). Therefore, sexual differentiation of PGCs is closely associated with distinct cell cycle regulation.

Although the initiation mechanism of mitotic arrest in male germ cells remains unclear, several factors have been identified as regulators of the cell cycle. Retinoblastoma 1 (RB1), a major cell cycle regulator involved in cell proliferation, apoptosis and cell differentiation in somatic cells (Zacksenhaus et al. 1996), and p38 mitogen-activated protein kinase (MAPK), which is activated by several cellular stresses (Ono and Han 2000), were previously suggested as regulators of mitotic arrest in male germ cells (Spiller et al. 2010; Ewen et al. 2010). However, the relationships among each cascade are unclear.

An evolutionarily conserved RNA-binding protein, NANOS2, was previously identified as an essential factor for male germ cell differentiation (Tsuda et al. 2003). NANOS2 protein is expressed in testes from E13.5, and promotes male-type gene expression such as DNMT3L and TDRD9 (Suzuki and Saga 2008; Suzuki et al. 2016). NANOS2 is also involved in cell cycle regulation. In the absence of NANOS2, male germ cells are arrested at G0 at E14.5 but resume the cell cycle at E15.5 (Suzuki and Saga 2008; Saba et al. 2014a). Therefore, NANOS2 was suggested to function in maintaining the arrested cell cycle.

NANOS2 interacts with another RNA-binding protein, DND1, and a deadenylation component, CNOT1, and localizes to processing bodies, by which the expression of recruited RNAs may be repressed (Suzuki et al. 2016, 2010, 2012; Parker and Sheth 2007; Decker and Parker 2012; Shimada et al. 2019). The possible target RNAs regulated by NANOS2 have been searched by several strategies, including NANOS2-IP to detect interacting RNAs, and expression change analyses using *Nanos2*-KO and *Nanos2*-overexpression. However, only two genes have been identified as bona fide NANOS2 targets (Kato et al. 2016; Zhou et al. 2015). One is *Dazl,* which is strongly expressed in germ cells just after colonizing the gonads in both males and females

(Seligman and Page 1998). DAZL was reported to act as a licensing factor for germ cells to initiate sexual differentiation (Gill et al. 2011). It is required for germ cells to acquire competence to respond to RA to enter meiotic prophase in ovaries, whereas its expression is repressed by NANOS2 once germ cells enter the testes. As mitotic activity is resumed and STRA8 is up-regulated in the absence of NANOS2, NANOS2-null germ cells were thought to be feminized. However, it is unclear whether the repression of DAZL by NANOS2 is responsible for the maintenance of mitotic arrest and repression of feminization.

To address the above question, we first conducted double KO of NANOS2 and DAZL, and found that DAZL is not the only factor responsible for cell cycle repression. To explore more factors responsible cell cycle regulation, we conducted single-cell RNA sequencing (scRNA-Seq) analyses, and found that NANOS2 functions not only to maintain the arrested cell cycle, but also to induce mitotic arrest.

## Results

### *Dazl* suppression by NANOS2 is not sufficient to repress mitotic resumption

In the previous study, the excess expression of DAZL, a target of NANOS2, led to the failure of male-type gene expression, meiosis initiation and resumption of mitosis.

Therefore, *Dazl* suppression by NANOS2 is expected to regulate cell differentiation and the cell cycle in male germ cells (Kato et al. 2016). To address this issue, we generated *Nanos2* and *Dazl* double knock-out (dKO) germ cells only after E12.5 by injecting tamoxifen at E12.5 (see Methods). If *Dazl* suppression by NANOS2 is the main driver of male-type gene expression and cell cycle arrest, the dKO should rescue the phenotype. We stained DNMT3L as a marker of male differentiation, STRA8 as a marker of meiosis initiation and Ki67 as a marker of active mitosis. Contrary to our expectation, DNMT3L expression was not detected and STRA8 was still positive in dKO germ cells (Fig. 1A), suggesting that the suppression of *Dazl* by NANOS2 is not sufficient to promote male germ cell differentiation. Importantly, a previous report demonstrated that excess DAZL expression increased the responsiveness of germ cells to RA (Kato et al. 2016). However, the dKO cells exhibited STRA8 expression, suggesting that DAZL is dispensable for germ cells to respond to RA at this point. In addition, dKO cells exhibited positive signals for Ki67 (Fig. 1B). After quantification, the proportion of mitotically active cells in the dKO was similar to that in *Nanos2*-KO (Fig. 1B). This suggests that there are other gene targets of NANOS2 besides *Dazl*.

**Figure 1.**
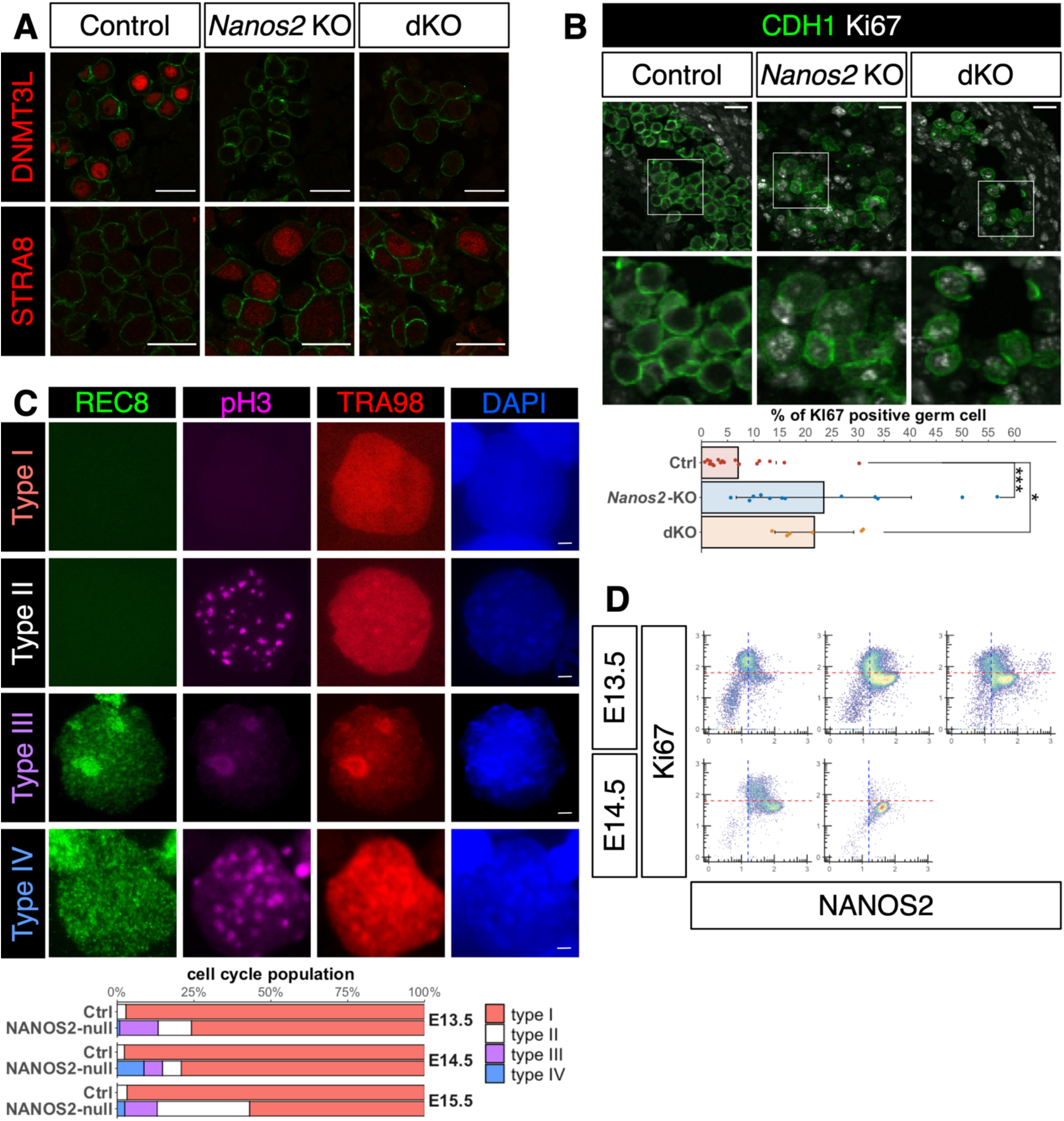
NANOS2 has functions separate from the suppression of *Dazl.* (A) Immunofluorescence analyses of testes from control, *Nanos2*-KO and *Dazl*; *Nanos2*-dKO at E15.5 using the male differentiation marker DNMT3L (red, upper) and meiosis marker STRA8 (red, lower). Tamoxifen was injected at E12.5. (B) Immunofluorescence analyses of testes from control, *Nanos2*-KO and *Dazl*; *Nanos2*-dKO at E15.5 using the mitosis marker (Ki67, white) and germ cell marker (CDH1, green). Magnified images in squares are also shown. The quantified data are shown below. ***P<0.001 and *P<0.05 (C) Classification of male germ cells (type I-IV) by immunofluorescence analysis with meiosis (REC8), mitosis (pH3) and germ cell (TRA98) markers. Quantification of control and NANOS2-null germ cells based on the classification at E13.5, E14.5 and E15.5. The bar graph is composed of type I (REC8^-^/pH3^-^: including mitotic arrest, orange), type II (REC8^-^/pH3^+^: mitosis, white), type III (REC8^+^/pH3^-^: meiosis, purple) and type IV (REC8^+^/pH3^+^: abnormal, blue). (D) Flow cytometric analysis of germ cells obtained from wild-type E13.5 testes. Cells were stained with anti-NANOS2 and anti-Ki67 antibodies. The blue dashed lines indicate the threshold of NANOS2 signal (see also Fig. S1 and S2). The red dashed lines indicate the threshold of Ki67 signal. The left side samples were speculated to be earlier stages based on NANOS2 expression. Scale bars: (A and B) 20 μm, (C) 1 μm.

### NANOS2 is required for the initiation of cell cycle arrest

To investigate the mechanism of cell cycle regulation by NANOS2, we re-examined cell cycle state changes during male germ cell development using *Nanos2*-hetero*^-^* and *Nanos2*-KO*^-^* testes from E13.5 to E15.5. The progression of mitosis and meiosis was examined in each single germ cell by co-staining with anti-phosphorylated histone H3 (pH3), a marker of M-phase, and anti-REC8, a meiosis specific cohesion, antibodies. In *Nanos2*^+/-^, we observed two types of cells exhibiting 1) negative signals for both REC8 and pH3 (type I), and 2) punctate signals only for pH3 (type II) (Fig.1C). The former cells likely included mitotically arrested cells, whereas the latter included proliferating cells because of the punctate pH3 signal (Barber et al. 2004). The majority of cells was classified as type I at E13.5 (97.1%), E14.5 (97.7%) and E15.5 (97.6%) (Fig. 1C), suggesting that mitotic arrest was initiated by E13.5.

We next examined NANOS2-null germ cells. In addition to type I and type II, NANOS2-null germ cells consisted of 3) REC8-single-positive cells, which most likely entered meiosis (type III), and 4) REC8 and pH3-double-positive cells, which were never observed among control XY or XX germ cells (type IV) (Fig. 1C). Quantification analysis revealed that 75.8% of germ cells were REC8/pH3-double-negative (type I) in NANOS2-null at E13.5 (Fig.1C). This proportion was retained at E14.5 (79.1%). However, the proportion of type I cells decreased to 56.5% at E15.5 and that of mitotically active type II cells increased (Fig. 1C). This demonstrated that mitotic cells increased at E15.5, consistent with previous observations, but a clear distinction was already observed at E13.5, suggesting that NANOS2 functions in both the maintenance and initiation of mitotic arrest.

To further examine cell cycle states, we stained germ cells with antibodies against NANOS2, TRA98 and Ki67, and analyzed them by fluorescence-activated cell sorting (FACS) at E13.5 and E14.5. Due to the variation in developmental timing, we observed heterogeneous results at these stages among samples (Fig. S1 and S2). Approximately 20% of cells were identified as germ cells in testes and the number of NANOS2-positive cells gradually increased with development (Fig. S1 and S2). Of note, NANOS2 protein was detected in the mitotic cells, confirming that NANOS2 expression starts prior to the downregulation of Ki67 (Fig. 1D). Once NANOS2 was expressed, Ki67 was strongly reduced, suggesting that NANOS2 is involved in the initiation of cell cycle arrest.

### Characterization of gonadal cell types and their sexual properties based on scRNA-Seq data

During embryonic germ cell development, the cell cycle status dynamically changes (Western et al. 2008; Miles et al. 2010). Upon sexual differentiation in particular, marked changes must be induced asynchronously in the transition period from E12.5 to E14.5. To analyze gene expression changes in the transition period and identify the initiation mechanism of mitotic cell cycle arrest, we analyzed the cell cycle state and gene expression profile at single-cell resolution.

We constructed single-cell RNA sequencing (scRNA-Seq) libraries using E11.5 to E15.5 testes and ovaries as the materials, and conducted transcriptome analyses. As an excess cell number reduces the sequence depth per cell, we omitted E14.5 female data from the following analysis (Fig.S3A and B). We used whole gonads as materials and our data contained information for numerous cell types. The t-SNE plot analysis separated at least 10 distinct clusters, C1 to C10 (Fig. 2A). To identify the cell type of each cluster, we examined representative gene expression in each cluster (Fig. S4).

**Figure 2.**
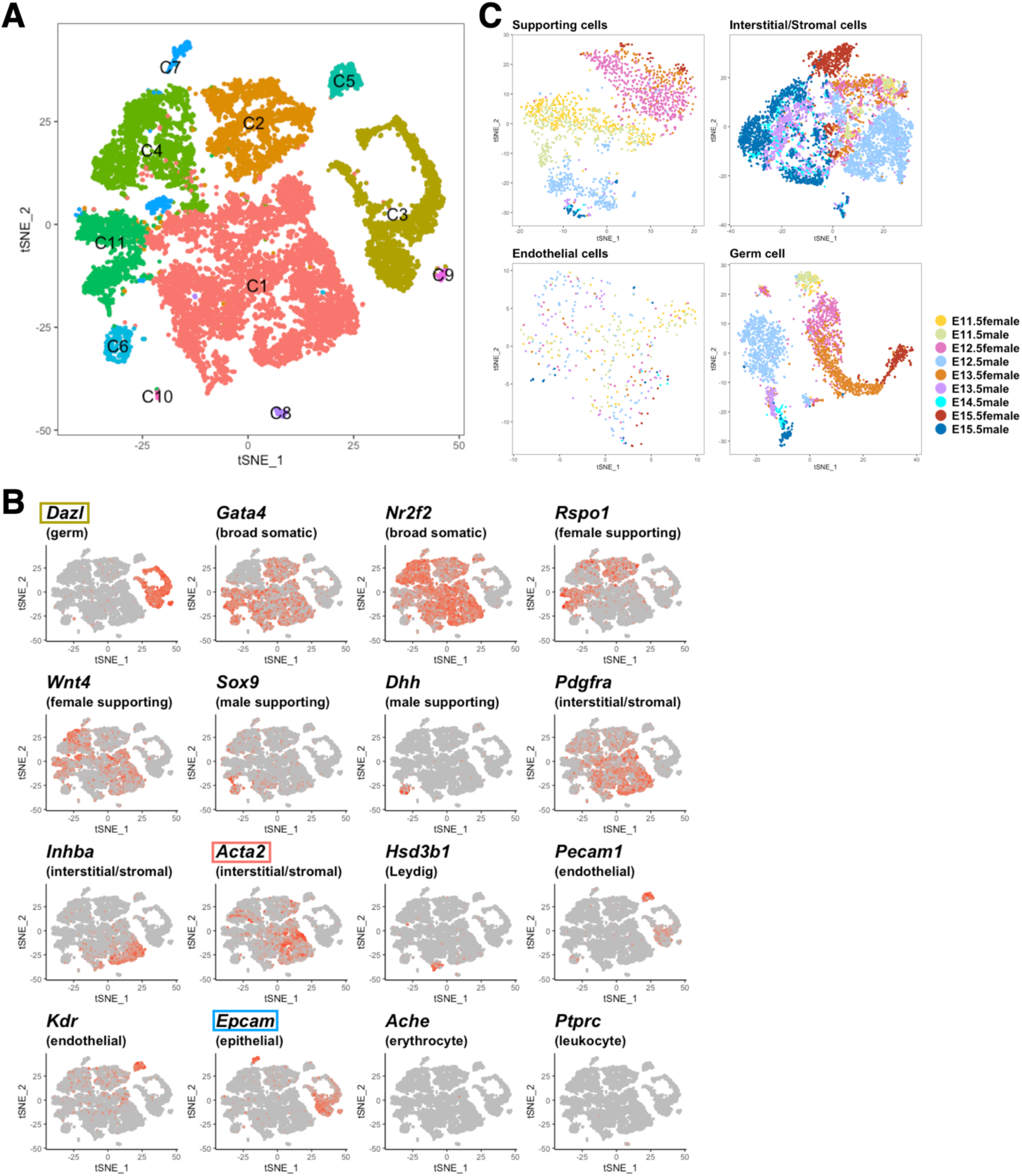
Cell type identification in scRNA-Seq data. (A) A tSNE plot for all cells. Each dot indicates each cell. Cells are colored for each cell cluster. (B) Expression levels of cell type marker genes are shown. Gene names are written in *italic* and cell types are indicated in parentheses. Cells with high expression are colored darker red. The gene names surrounded by rectangles are shown in the heat map of Fig. S4. (C) tSNE plots for supporting, interstitial/stromal, endothelial or germ cells. Cells are colored by embryonic stage.

Several representative specific cell type markers, such as *Acta2* in C1, *Dppa5a*, *Dppa3* and *Dazl* in C3, *Aldh1a1* in C6, *Epcam* in C7, *C1q* genes in C8, *Rhox* genes in both C3 and C9, *hemoglobin* genes in C10 and *Fst* in C11, were listed. In addition, many novel genes exhibiting certain cell population-specific expression were observed, demonstrating our scRNA-Seq data to be useful to analyze the cell population in developing gonads. Using known distinct genes as markers, we successfully identified the germ cell population and other somatic cell populations (Fig. 2B). Importantly, we were able to observe the heterogeneity of gene expression within each cluster, especially in C1, suggesting that more careful analysis can separate more detailed cell types. To evaluate whether our data can be used to analyze detailed differentiation pathways, such as sexual differentiation, we extracted 4 major cell types in the gonad; germ, supporting, interstitial/stromal and endothelial cell populations, and performed clustering analysis for each separately (Fig. 2C). Based on the tSNE plot, some of the E11.5 male supporting cells were already separated from female cells and located closer to E12.5 male cells (Fig. 2C), indicating that male-type differentiation already started via SRY. Although *Sry* itself was not detected in our scRNA-Seq data, downstream genes of SRY, *Sox9* and *Fgf9*, were activated in these cells (Fig. S3C). The interstitial/stromal cell population at later stages (E14.5 and E15.5) was clustered based on sex, but they were still clustered in a similar position at E12.5, suggesting that their sexual features were determined after the sexual differentiation of supporting cells. The endothelial cells did not exhibit clear segregation based on sex, suggesting that they do not have a clear sexual difference within this time window. At E11.5, germ cells from testes (male, light green) and ovaries (female, light yellow) had almost identical properties, but after E12.5, they exhibited a clear sex-specific differentiation trajectory (Fig. 2C). These data suggest that sex determination occurs in supporting cells at E11.5 and is followed by the sexual differentiation of other cell types, including germ cells. This observation is consistent with the previous reports that SRY activates SOX9 expression in supporting cells (Sekido et al. 2004) and these male supporting cells promote the male-type differentiation of surrounding cells by secreting FGF9 (Kim et al. 2006; Colvin et al. 2001). Therefore, we concluded our scRNA-Seq data to be sufficiently reliable to analyze the sexual differentiation of each cell type.

### Cell cycle changes during the differentiation of germ cells

We extracted the germ cell population and conducted further analyses (Fig. 2C). To characterize the cell cycle state of each cell, we conducted cell cycle analysis using a previously reported method (Macosko et al. 2015; Kashima et al. 2018). As germ cells have a specific cell cycle phase, meiosis, we further modified the method to apply to germ cells (see Methods for details). This method enabled the visualization of each cell cycle state during germ cell differentiation. As expected, most germ cells were mitotically active at early stages (E11.5 and E12.5) (Fig. 3A). After segregation into a sex-specific direction, cell cycle states demonstrated sex specificity. In the female trajectory, meiotic cells accumulated from E13.5 and reached almost 100% at E15.5. In the male trajectory, mitotically arrested cells accumulated from E13.5 and most cells entered the arrested state at E14.5 (Fig. 3A and B). This is consistent with previous reports (Western et al. 2008; Miles et al. 2010), supporting the use of our scRNA-Seq data to analyze the cell cycle regulation during germ cell development.

**Figure 3.**
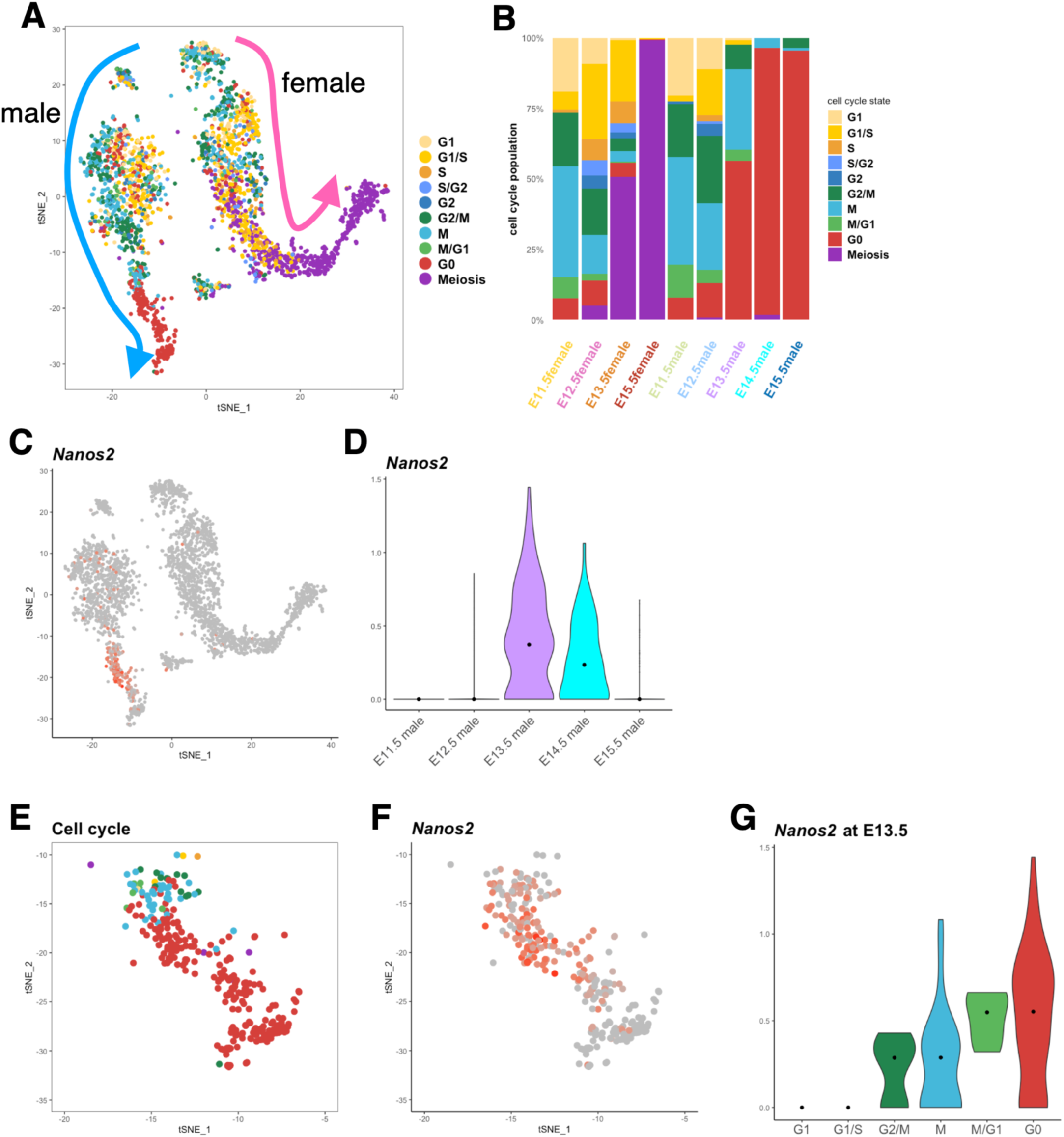
Relationship between cell cycle arrest and *Nanos2* expression. (A-B) Cell cycle estimation of germ cells. (A) shows the tSNE plot for germ cells. Cells are colored by the estimated cell cycle phase. Differentiation directions estimated by embryonic stage and sex of each cell (refer to Fig. 2C) are indicated as arrows. The quantified data are shown in (B). (C) Expression levels of *Nanos2* are shown. Cells with high expression are colored darker red. (D) Violin plot for *Nanos2* expression. Black dots indicate the median in each stage. € An enlarged tSNE plot to focus on E13.5-E15.5 male germ cells. Cells are colored by the estimated cell cycle. (F) Expression levels of *Nanos2* are visualized on the tSNE plot of (E). (G) Violin plot for *Nanos2* expression at E13.5 in each cell cycle phase. Black dots indicate the median in each phase.

### *Nanos2* expression precedes cell cycle arrest

As shown in Fig. 1F, NANOS2 expression precedes mitotic arrest. Using scRNA-Seq data, we examined the relationship between *Nanos2* expression and the cell cycle in detail. *Nanos2* expression was detected only in male germ cells at E13.5 and E14.5 (Fig. 3C and D). E13.5 is the timepoint when the cells enter G0 arrest (Fig. 3A and B). To correlate *Nanos2* expression and cell cycle status, we measured *Nanos2* expression levels in each cell cycle state at E13.5 (Fig. 3E-G). We found that *Nanos2* expression started in cells in M-phase at E13.5, and when *Nanos2* expression peaked, the G0 population accumulated (Fig. 3G). These observations support the idea that NANOS2 functions to induce cell cycle arrest.

### Cell cycle regulation in NANOS2-null cells

If NANOS2 is required for the induction of cell cycle arrest, NANOS2-null cells should fail to enter mitotic arrest. To examine this, we also performed scRNA-Seq using *Nanos2*-KO testes at E13.5 and E14.5, and analyzed these germ cell data together with wild-type data. As NANOS2-null cells exhibited meiotic marker gene expression, such as STRA8 and SYCP3 (Suzuki and Saga 2008), which are widely used as female markers at embryonic stages, we expected NANOS2-null male germ cells to be plotted to closer to the wild-type female trajectory. However, NANOS2-null cells were plotted to the male side, indicating that these cells retained male properties (Fig. 4A). RNA velocity analysis (La Manno et al. 2018), which enables the visualization of the differentiation direction and strength, also demonstrated that NANOS2-null cells failed to progress to male-type differentiation but were not feminized (Fig. 4B).

**Figure 4.**
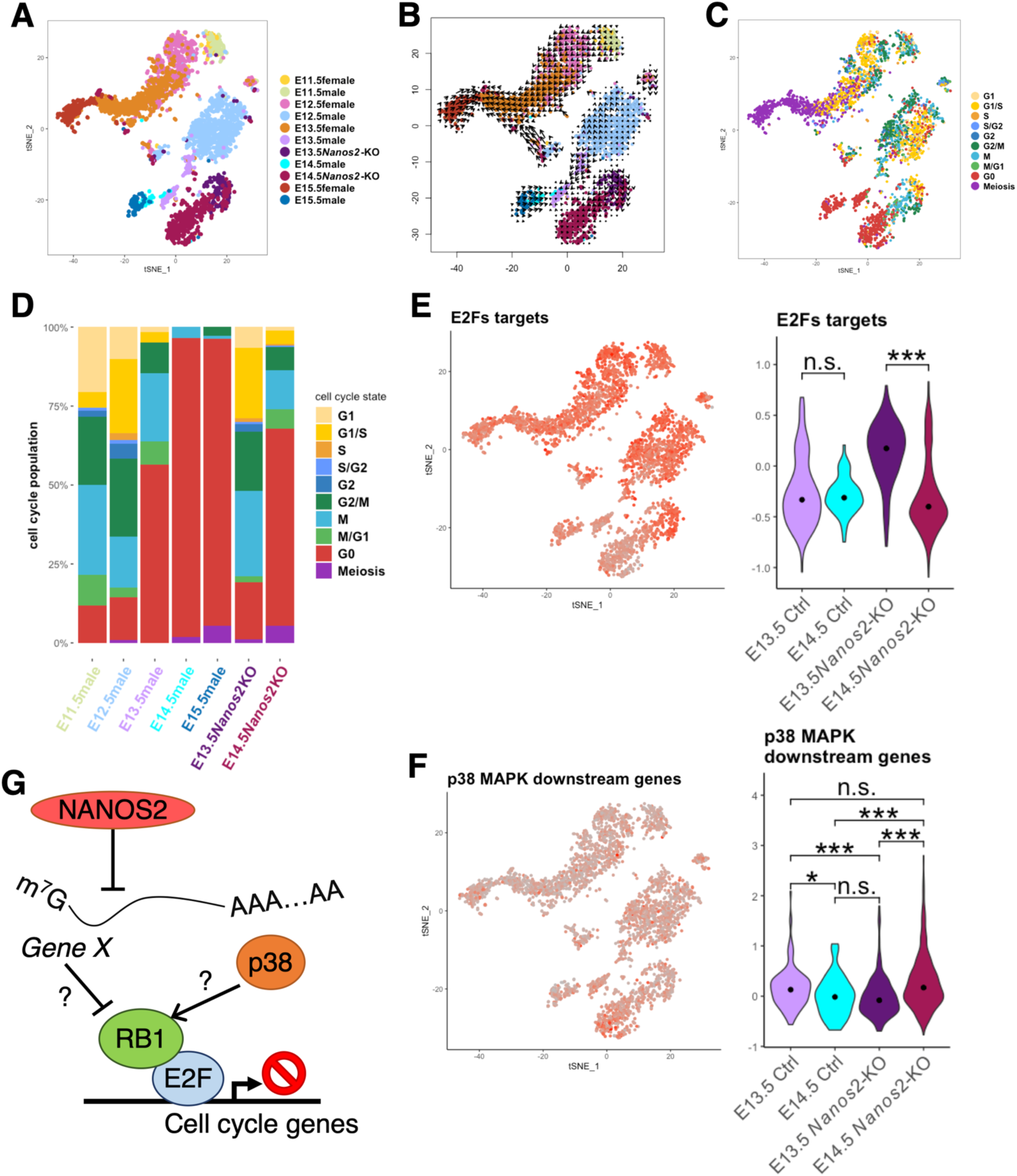
NANOS2 is required for the initiation of cell cycle arrest. (A-D) tSNE plot for wild-type and *Nanos2*-KO germ cells. Cells are colored by embryonic stage and genotype in (A), and cell cycle phase in (C). The quantified data of (C) are shown in (D). (B) RNA velocity was projected on the tSNE plot. Arrows indicate directions of cell differentiation, and the strength was estimated by the expression profile of spliced and un-spliced forms of RNAs. (E and F) Average expression values of E2Fs target genes (E) and downstream genes of p38 MAPK (F). *Cdc25*, *Mcm2-7*, *Fen1*, *Kif11*, *Smc2*, *Smc4*, *Ect2*, *Ccne2*, *Cenpe*, *Cdca8*, *Hmgb2*, *Bub1*, *Kif20a*, *Ndc80*, *Kif18a*, *Ncapg*, *Ccna2* and *Nekl2* were used to calculate the E2Fs activity, and *Mapkapk5*, *Max*, *Myc*, *Hbp1* and *Cebpa* were used to calculate the p38 MAPK activity. Expression values are visualized on tSNE plots (left) and violin plots (right). Black dots indicate the median. ***P<0.001, **P<0.01 and *P<0.05 (G) A scheme of the proposed mechanism of cell cycle regulation in male germ cells.

Cell cycle analysis revealed that most NANOS2-null germ cells were mitotically active at E13.5, similar to wild-type male germ cells at E12.5 (Fig. 4C and D). This suggests that NANOS2-null germ cells failed to enter the arrested stage at E13.5. However, approximately 70% of NANOS2-null cells were arrested at the G0 stage at E14.5 (Fig. 4D), being consistent with a previous report (Suzuki and Saga 2008; Saba et al. 2014a) and implying that NANOS2-independent mechanisms to repress cell cycle progression also function in male germ cells. Consistent with these cell cycle changes in NANOS2-null germ cells, Gene Enrichment analysis demonstrated that cell cycle-related genes were highly upregulated in NANOS2-null cells at E13.5; however, these terms were not listed at E14.5 (Fig. S5).

To identify the NANOS2-independent pathway, we examined reported cell cycle regulators implicated in male germ cells other than NANOS2 such as Retinoblastoma 1 (RB1) and p38 MAPK (Spiller et al. 2010; Ewen et al. 2010). RB1 represses cell cycle progression by suppressing E2F function, which activates S phase genes (Wu et al. 1995; Kent and Leone 2019). p38 MAPK is activated by numerous extracellular stimuli and is considered to lead to mitotic arrest in male germ cells (Ewen et al. 2010). Moreover, p38 MAPK regulates the cell cycle by promoting RB1 function (Gubern et al. 2016). To visualize the activity of each signal, we calculated the average level of scaled known target gene expression values and visualized them in tSNE and violin plots (see Methods for detail). In a previous report, signals for the phosphorylated inactive form of RB1 (pRB1) were low at E14.5 and increased at E15.5 in the NANOS2-null germ cells (Saba et al. 2014a). Consistent with this, the averaged level of E2Fs-target gene expression was high at E13.5 but decreased at E14.5 in *Nanos2*-KO samples (Fig. 4E and S6). This correlated well with the changes in cell cycle states from E13.5 to E14.5 in NANOS2-null germ cells, suggesting that both NANOS2-dependent and -independent mechanisms regulate the cell cycle via RB1 function. Because p38 are activated only in male, we selected 5 genes which showed male enriched expression to visualize the p38 MAPK activity (Ewen et al. 2010). Then, we found that p38 MAPK activity slightly increased in E14.5 NANOS2-null germ cells (Fig. 4F and S7A), suggesting p38 MAPK as a possible candidate to suppress the cell cycle in NANOS2-null germ cells at E14.5. We also confirmed that mRNA expressions of p38 MAPK genes were increased at E13.5 but not at E14.5 in NANOS2-null cells (Fig. S7B). This implies that the increase of p38 MAPK activity is mainly caused by the hyperactivation. Taken together, we propose that NANOS2 and p38 MAPK work in parallel to suppress the cell cycle by regulating RB1 function (Fig. 4G).

### Identification of targets of NANOS2 to regulate the cell cycle

As the primary key function of NANOS2 may be to initiate mitotic arrest in male germ cells, it is important to identify NANOS2 targets that achieve this function. To detect possible direct targets, we focused on the expression changes in spliced forms of transcripts in NANOS2-null germ cells because NANOS2 functions in the cytoplasm and acts on only spliced RNAs (Fig. 5A). For this purpose, we examined gene expression changes in spliced and un-spliced forms separately using our scRNA-Seq data with the velocyto package (see Materials and Methods for details) (La Manno et al. 2018). We first identified differentially expressed genes (DEGs) in their spliced forms in NANOS2-null male germ cells at E13.5 and E14.5, and extracted 834 and 1,678 DEGs, respectively (table S1 and S2). This population also contained transcriptionally regulated genes. To obtain only post-transcriptionally regulated genes, we selected genes merged with the transcriptionally unchanged population (unDEGs in un-spliced form) (Table S3). We identified 743 and 1,358 gene candidates at E13.5 and E14.5, respectively, and 319 genes were commonly listed in both stages (Fig. 5B). The DEGs contained 187 and 111 commonly up- or down-regulated genes, respectively, in the NANOS2-null germ cells between E13.5 and E14.5 (Fig.5C).

**Figure 5.**
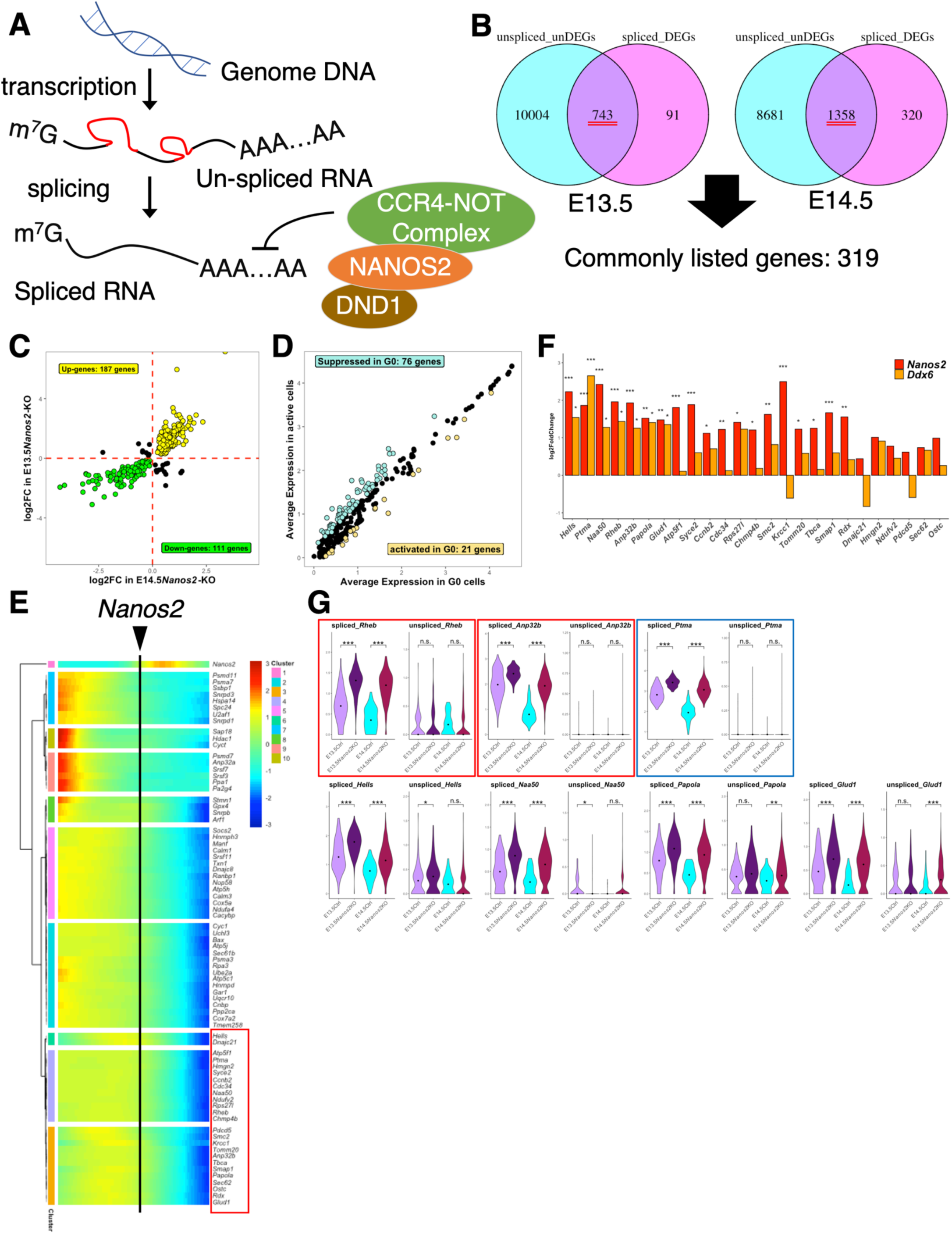
Identification of target gene candidates suppressed by NANOS2. (A) The scheme of the mRNA-suppressing mechanism by NANOS2. NANOS2 makes a complex with DND1 and the CCR4-NOT complex via CNOT1, and suppresses spliced mature RNAs. (B) Venn diagrams comparing unDEGs of un-spliced forms and DEGs of spliced forms in NANOS2-null germ cells at E13.5 (left) and E14.5 (right). In total, 743 and 1,358 genes were listed as spliced form-specific DEGs in E13.5 and E14.5, respectively, and 319 genes were commonly listed. (C) The scatter plot to compare the log2 fold change in *Nanos2*-KO samples at E13.5 (y axis) and E14.5 (x axis). Upregulated genes in the KO both at E13.5 and E14.5 (Up-genes) are colored in yellow and downregulated genes in the KO (Down-genes) are colored in green. Red dashed lines are drawn at 0. (D) The gene expression of 187 up-regulated genes in G0 arrest (x axis) and the active cell cycle (y axis) in male germ cells at E13.5 are shown. Average expression levels were calculated for each condition and plotted. Light blue dots were suppressed and orange dots were activated in G0 cells. (E) The gene expression patterns of the 76 obtained gene candidates suppressed by NANOS2 along the developmental pseudo-time of male germ cells. The point when *Nanos2* expression became strong is indicated as a vertical line. Genes surrounded by a red rectangle belong to a gene group exhibiting downregulation just after *Nanos2* expression. (F) The expression changes in identified genes in (E) in *Nanos2*-KO (red) and *Ddx6*-KO (orange) male germ cells at E16.5. ***P<0.001, **P<0.01 and *P<0.05. (G) The expression changes in identified genes in (F). Genes surrounded by rectangles, *Rheb*, *Anp32b* and *Ptma,* showed significant changes only in the spliced form at E13.5 and E14.5. Red rectangles indicate mTORC1-related genes (see Fig. 6 for details).

To identify gene candidates involved in cell cycle arrest, we searched for genes exhibiting expression difference between G0 arrested and mitotically active cells at E13.5 in the wild-type male. Among 187 up-regulated genes in NANOS2-null germ cells, 76 were suppressed in G0 cells, and among 111 down-regulated genes, 21 were activated in G0 cells (Fig. 5D).

We reasoned that NANOS2 target genes exhibit correlated gene expression changes with *Nanos2* expression during male germ cell differentiation. Thus, we conducted pseudo-time analysis for the obtained gene candidates (Trapnell et al. 2014; Qiu et al. 2017a). In this analysis, male germ cells were aligned to make a single-directional trajectory. Therefore, gene expression changes along cell differentiation can be visualized (Fig. 5E). By selecting genes exhibiting expression changes just after *Nanos2* expression, we obtained 25 and 19 genes as candidates being suppressed and activated by NANOS2, respectively (Fig. 5E and S8). As the main function of NANOS2 is the suppression of target genes, we further focused on the 25 suppressed candidates.

We previously reported that NANOS2 regulates the cell cycle in a DDX6-dependent manner (Shimada et al. 2019). Therefore, we expected commonly up-regulated genes in both NANOS2-null and DDX6-null germ cells to be strong candidates. By these criteria, we finally obtained 7 genes, *Hells*, *Ptma*, *Naa50*, *RHeb*, *Anp32b*, *Papola* and *Glud1* (Fig. 5F). Among them, *Rheb*, *Anp32b* and *Ptma* were confirmed to be up-regulated only in spliced form in the NANOS2-null germ cells (Fig. 5G). Importantly, all 3 genes are known to be involved in cell proliferation (Bianco and Montano 2002; Vareli et al. 1996; Sun et al. 2001; Yang et al. 2016; Hao et al. 2018; Sancak et al. 2008). Therefore, we concluded that these 3 genes are possible NANOS2 target candidates to regulate the cell cycle.

### NANOS2 directly suppresses target candidates in cultured cells

NANOS2 was reported to repress mTORC1 activity in spermatogonial stem cells (Zhou et al. 2015) and *Rheb* is an activator of mTORC1 (Sancak et al. 2008; Hao et al. 2018). In addition, Akt can activate mTORC1 via the inhibition of TSC function and ANP32B is known to activate Akt (Skeen et al. 2006; Inoki et al. 2002; Yang et al. 2016) (Fig. 6A). Therefore, we expected NANOS2 to suppress mTORC1 activity via the suppression of *Rheb* and *Anp32b*. To address this possibility, we examined mTORC1 activity using phosphorylated ribosomal protein S6 signals (pS6), a key output of mTORC1 signaling. In control testis, no germ cells exhibited positive signals for pS6 at E15.5. In contrast, approximately 9% of NANOS2-null germ cells had strong pS6 signals (Fig. 6B), suggesting that mTORC1 signaling was suppressed under the control of NANOS2.

**Figure 6.**
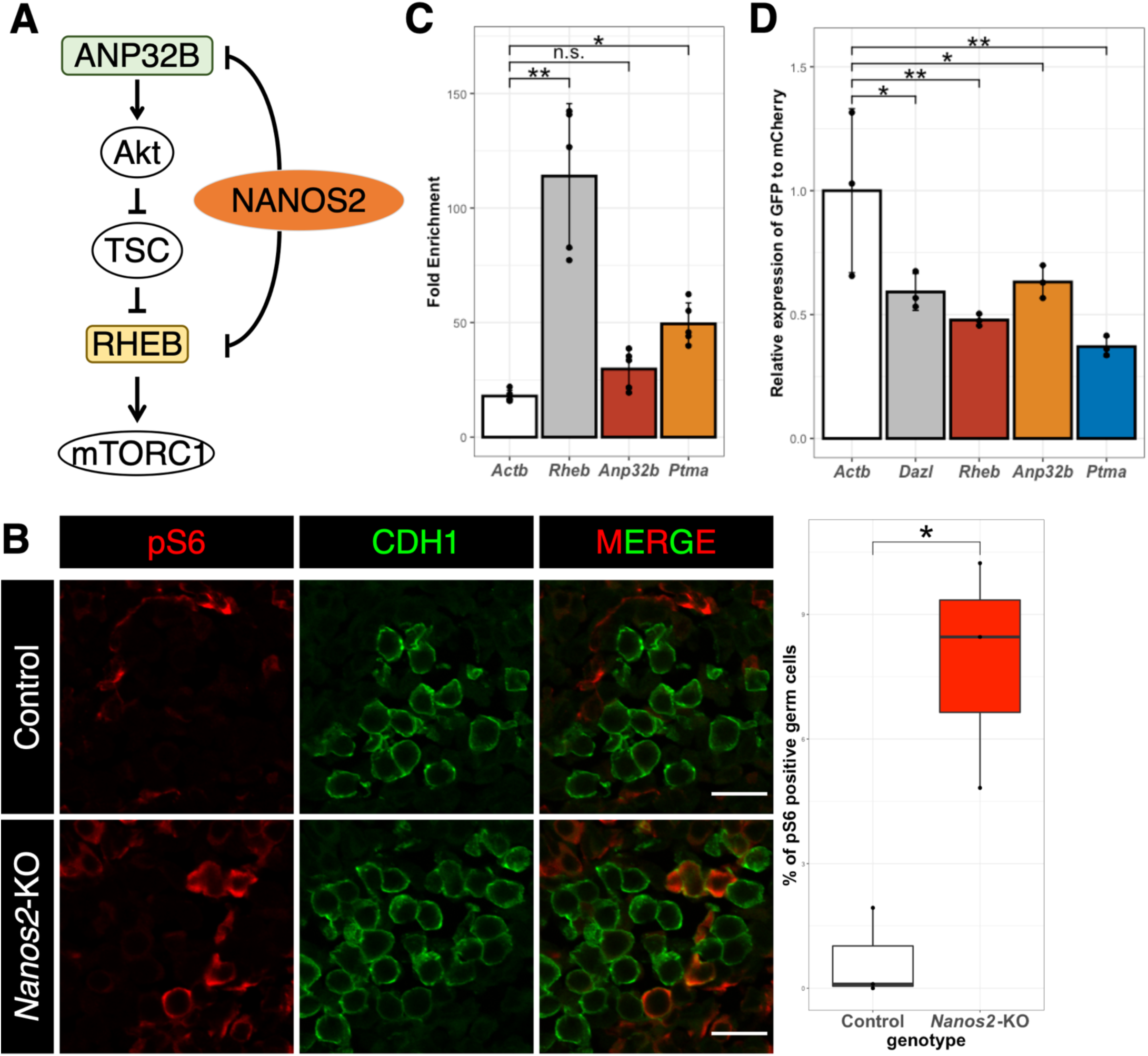
NANOS2 suppresses mTORC1 activity via the suppression of its activator. (A) A scheme of the regulatory pathway for mTORC1 activation. ANP32B and RHEB can activate mTORC1, and their mRNA expression is suppressed under the control of NANOS2 in male germ cells. (B) Immunofluorescence analyses of testes from control and *Nanos2*-KO at E15.5 using the active mTORC1 marker pS6 (red) and germ cell marker CDH1 (green). Quantified data are shown on the right. Three independently prepared samples were analyzed. *P<0.05. (C) RNA-IP experiments conducted using the cell culture system (see Methods for details). The endogenous mRNAs were detected using RT-qPCR. n=3. **P<0.01, *P<0.05 and n.s.=not significant. (D) The relative expression value of *Emgfp* is shown. The *Emgfp* value was normalized by the *mCherry* value, and the relative value to the *Actb* 3’UTR was calculated. **P<0.01 and *P<0.05.

To assess whether NANOS2 directly regulates these candidate mRNAs, we conducted RNA-IP analysis. As the germ cell number is limited in the testes, we utilized a cultured cell system developed in our laboratory in which cell cycle arrest can be induced upon doxycycline (Dox)-inducible NANOS2 and DND1 expression (Hirano et al., manuscript in preparation). In this system, 3xFlag-tagged NANOS2 and its essential partner, HA-tagged DND1, can be induced by the addition of Dox, and NANOS2-target RNAs should be co-precipitated using anti-FLAG antibody. As these target gene candidates are ubiquitously expressed in both germ and somatic cells (Fig. S9), we reasoned that NANOS2 expressed in cultured cells can interact with endogenously expressed mRNA. Co-precipitated RNAs were quantified by RT-qPCR. Compared with the negative control *Actb*, *Rheb* and *Ptma* were enriched in the IP sample (Fig. 6C), indicating that *Rheb* and *Ptma* are possible direct targets of NANOS2.

Lastly, to examine the effects of NANOS2 on the bound RNAs, we conducted NANOS2 functional analyses using this cultured cell system. We previously reported that NANOS2 regulates *Dazl* expression by its binding to the 3’UTR (Kato et al. 2016). Thus, we expected these candidates to also be regulated through the binding of NANOS2 to their 3’UTR. We cloned the 3’UTR of gene candidates into a dual-fluorescence reporter vector (Fig. S10A and B), in which mCherry serves as an internal control and EmGFP reports the activity of the 3’UTR of target RNAs. In the absence of NANOS2 and DND1 (DMSO treatment), EmGFP containing the candidate 3’UTR did not exhibit significant expression reduction compared with *Actb* control 3’UTR (Fig. 10C). In contrast, in Dox-treated cells, all constructs exhibited significant downregulation comparable to the repression level of the *Dazl* 3’UTR (Fig. 6D), suggesting that NANOS2 suppresses the expression of these target genes in a 3’UTR-dependent manner.

## Discussion

Arrest of the cell cycle is known as the first step of male-type differentiation in germ cell. It was previously suggested that NANOS2 represses *Dazl* for the maintenance of cell cycle arrest by regulating RA responsiveness (Kato et al. 2016). However, we revealed that *Dazl* is not the only target of NANOS2 to stop the cell cycle and the scRNA-seq confirmed that NANOS2 played a role in both the maintenance and initiation of cell cycle arrest. In this regard, we identified additional targets, *Rheb* and *Anp32b,* which function in the mTORC pathway separately from DAZL (Fig. 7). Although *Anp32b* was not significantly enriched in the IP sample, it exhibited spliced form-specific expression changes in NANOS2-null cells (Fig. 5 and 6) and 3’UTR-dependent downregulation by NANOS2, implying that it is not a direct target but still acts under the control of NANOS2 (Fig. 6D). NANOS2 was reported to suppress mTORC1 activity by anchoring MTOR protein in processing bodies in spermatogonial stem cells (Zhou et al. 2015). This report and our current data suggest that NANOS2 can regulate mTORC1 activity in multiple ways. However, we observed positivity of the active mTORC1 marker pS6 in some NANOS2-null germ cells. This suggests that there is another pathway to mediate the effects of NANOS2. As one candidate, we identified *Ptma* as a NANOS2 target. PTMA is ubiquitously expressed in mice (Eschenfeldt and Berger 1986) and its expression is regulated by E2Fs. Inhibition of *Ptma* by antisense oligomer prevents cell division (Sburlati et al. 1991). As mTORC1 activity promotes Cyclin D1 protein expression (Averous et al. 2008) and Cyclin D1 is known to activate E2F function via the inhibition of RB1 (Resnitzky and Reed 1995), *Ptma* may be suppressed both transcriptionally and post-transcriptionally under the control of NANOS2. PTMA expression peaks at around the late S or G2 phase; therefore, it is expected to function at these cell cycle phases. Taken together, NANOS2 may regulate different cell cycle regulatory machineries at different cell cycle phases to effectively suppress the cell cycle.

**Figure 7.**
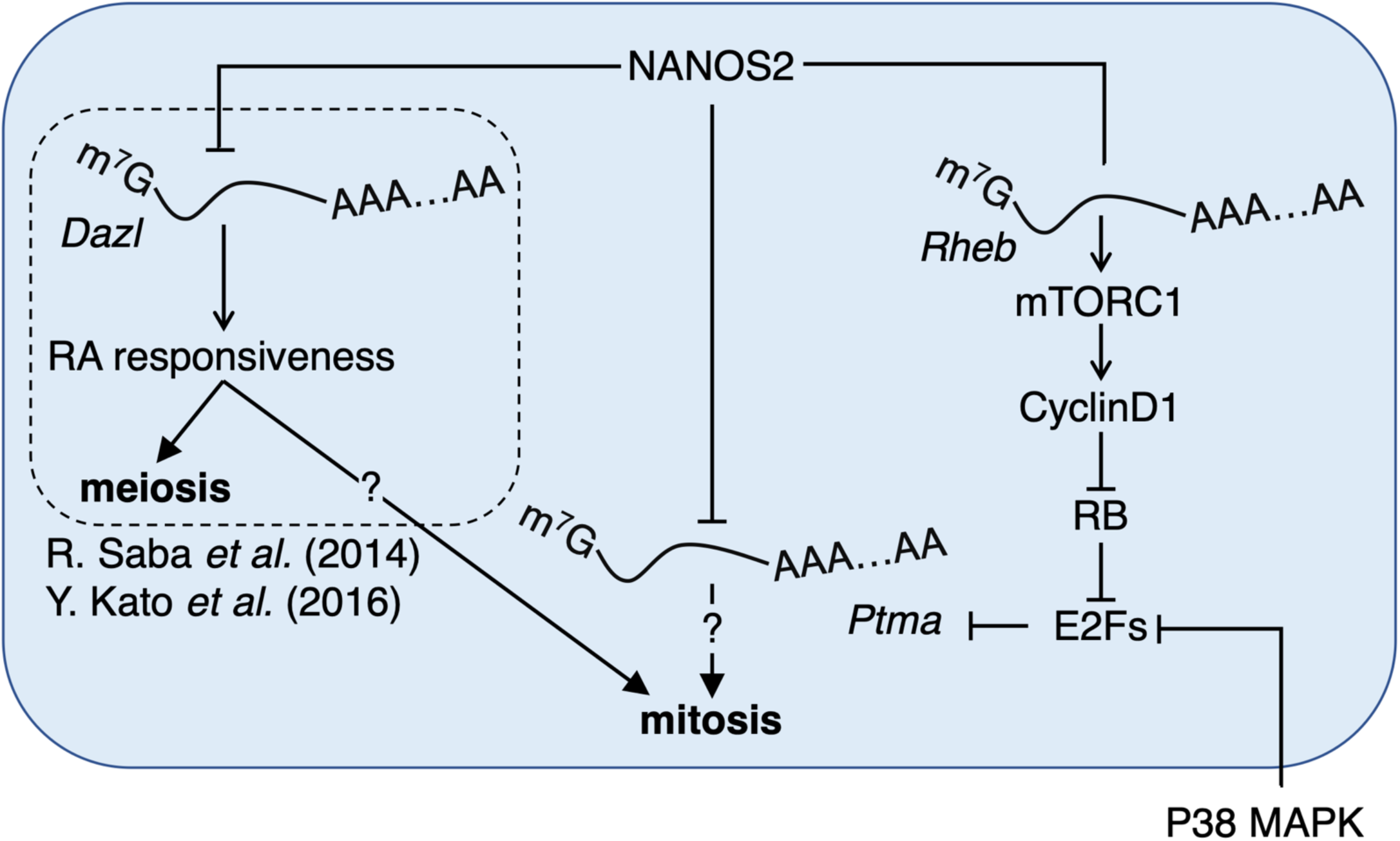
Gene network for cell cycle regulation during male germ cell development. Schematic drawing of the model proposed in this and previous studies. In the previous study, *Dazl* was identified as the direct target of NANOS2 and DAZL may increase RA responsiveness. Moreover, RA signaling plays an important role in cell cycle regulation in germ cells. We demonstrated that a more complex gene network exists under the control of NANSO2. NANOS2 downregulates mTORC1 activity via the suppression of its activator(s). This results in the downregulation of CyclinD1 and activates the RB1-suppressing E2F function. As an independent pathway, NANSO2 also suppresses *Ptma* and PTMA translation is suppressed by the suppression of mTORC1 activity. p38 MAPK may also function in parallel with NANOS2 to suppress RB.

In addition to cell cycle regulation, NANOS2 represses meiotic initiation by repressing STRA8, keeps germ cells in the seminiferous tubules and, importantly, promotes male-type gene expression (Saba et al. 2014a; Suzuki and Saga 2008). We believe that other NANOS2 targets contribute to these important events. Based on our pseudo-time expression data, many other genes are down-regulated just after *Nanos2* expression. However, there may not be a single responsible gene, as demonstrated for cell cycle regulation, which makes further functional analyses difficult.

Apart from NANOS2 function, our scRNA-Seq data are useful to analyze the sex determination mechanism of germ cells occurring between E11.5 and E12.5 in male gonads. Although the genetic cascades leading to the sexual differentiation of somatic cells are well studied, somatic factors to induce male germ cell determination have not been identified. As our data include whole gonad information and we confirmed that sexual differentiation starts earlier in supporting cells than in germ cells (Fig. 2C), further studies focusing on the signal transduction pathways are expected to find causative reciprocal interactions.

Although we focused only on genes repressed by NANOS2 associated with cell cycle arrest in this study, scRNA-Seq identified many genes that were up-regulated following NANOS2 expression. As the major function of NANOS2 is considered to be the repression of gene expression, such up-regulation may be a secondary effect. Further functional analyses will delineate a gene network by which male germ cell differentiation is orchestrated.

## Materials and Methods

### Mice

All mouse experiments were approved by the National Institute of Genetics (NIG) Institutional Animal Care and Use Committee. Mice were housed in a specific pathogen-free animal care facility at NIG. MCH (provided by CLEA Japan, Inc.), *Nanos2* KO (Tsuda et al. 2003), *Dazl* conditional KO (Fukuda et al. 2018), and *Oct4dPE-*CreER^T2^ (provided by A. Suzuki, Yokohama National University) mice lines were used in this study. To generate *Nanos2* and *Dazl* dKO mice, *Nanos2*-KO mice were crossed with *Dazl* conditional KO and *Oct4dPE*-CreER^T2^-carrying mice, and 500 μl of 10 mg/ml tamoxifen (Sigma-Aldrich, T5648) was intraperitoneally injected to knock-out *Dazl*.

### Preparation of germ cell spreads

Germ cell spreads were prepared from E13.5 to E15.5 testes. Testes were collected in 700 μl of M2 medium, passed through an 18-G needle and washed with PBS. The samples were incubated with 1 mg/ml of collagenase Type I (Invitrogen) at 37°C for 15-30 min, washed with PBS and incubated with Trypsin-EDTA for 1 min at 37°C. After washing with 500 μl of DMEM containing 10% (v/v) FCS and with PBS, hypotonic buffer (30 mM Tris-HCl (pH 7.5), 50 mM sucrose, 17 mM trisodium citrate dehydrate, 5 mM EDTA, 0.5 mM DTT and 0.5 mM PMSF) was added and incubated for 30 min at room temperature (RT). After spinning for 3 min at 400 × g, cells were resuspended in 100 mM sucrose containing 1 mM sodium hydroxide. The cell suspension was spread on slide glasses with 1% PFA containing 0.5% Triton X-100. Slide glasses were placed in a humidified slide chamber and incubated at 37°C overnight and then subjected to the immunostaining procedure. The spread germ cells were reacted with primary antibodies overnight at 4°C at the following dilutions: 1:100 for rabbit anti-REC8 (a gift from Dr. Watanabe, Tokyo University) and 1:200 for mouse anti-phospho-histone H3 (ser10) (a gift from Dr. Kimura, Osaka University). Secondary antibodies labelled with Alexa Fluor 488, 594 or 647 (1:1,000, Invitrogen) were used. DNA was counter-stained with DAPI (100 ng/ml). Fluorescence microscopy was performed using Olympus FV1200.

### Gonad sample preparation and immunostaining

Embryonic gonads were fixed in 4% PFA for 30 min at 4°C. Gonads were next submerged in 10 and 20% sucrose in PBS for 1 h each at 4°C, and in 30% sucrose in PBS overnight at 4°C. The gonads were then embedded in Tissue-Tek O.C.T. compound (Sakura Finetek, Tokyo, Japan) and frozen in liquid nitrogen. Six-micrometer-thick sections of each gonad were applied to glass slides and autoclaved in TRS (Dako). After pre-incubation with 3% skim milk in PBS-T (PBS with 0.1% Tween20 (Sigma-Aldrich)) at RT for 30 min, the sections were reacted with primary antibodies overnight at 4°C at the following dilutions: 1:200 for mouse anti-phospho-histone H3 (ser10) (provided by Dr. Kimura, Osaka University), goat anti-CDH1 (1μg/ml, R&D), rabbit anti-DNMT3L (1:500, provided by Dr. Yamanaka), rabbit anti-STRA8 (1:200, abcam#ab49602), rabbit anti-Ki67 (1:200, Invitrogen #MA5-14520) and anti-pS6 (1:200, CST #2211). Secondary antibodies labeled with Alexa 488, 594 or 647 (1:1,000, Invitrogen) were used. DNA was counter-stained with DAPI (100 ng/ml). Fluorescence microscopy was performed using Olympus FV1200 and images were processed with ImageJ.

### Flow Cytometry

Single-cell suspensions were prepared from testes from E13.5 and E14.5 embryos by incubation with 0.15% trypsin-EDTA at 37°C for 5 min, and then washed with DMEM containing 0.5% BSA (wash medium). After discarding supernatant, cells were fixed in 100 μl of 4% PFA for 10 min at RT, followed by the addition of 900 μl of cold 100% EtOH and incubated at 4°C overnight. After spinning down for 5 min at 400 × g, cells were washed with wash medium and reacted with primary antibodies for 1 h at RT at the following dilutions: mouse anti-NANOS2 (1:200), rabbit anti-Ki67 (Invitrogen #MA5-14520) and rat anti-TRA98 (1:4,000, a gift from Y. Nishimune, Osaka University). Then, cells were washed with wash medium and reacted with secondary antibodies labeled with Alexa Fluor 488, 594 or 647 (1:1,000, Invitrogen) for 30 min at RT. Then, washed cells were re-suspended in 0.1% BSA-containing PBS and analyzed by a JSAN Desktop Cell Sorter.

### Single-cell RNA-Seq

Single-cell suspensions were prepared from 6-26 gonads from E11.5-E15.5 embryos by incubation with 0.15% trypsin-EDTA at 37°C for 5-10 min. The E11.5 embryos were genotyped to determine the sex using primers to detect *Ube1,* as described in a previous report (Chuma and Nakatsuji 2001). After incubation, wash medium was added to inhibit trypsin and cells were filtered through a 35-μm sieve (BD Bioscience). Cells were re-suspended in DMEM containing 10% FBS and the cell density was adjusted to 1.0×10^6^ cells/ml. We tried to load 3,000 cells on the Chromium™ Controller (10x Genomics Inc.) and constructed the scRNA-Seq libraries using Chromium™ Single Cell 3’ Reagent Kits v2 following the manufacturer’s instructions. Each library was read at a depth of 1 million per sample for cell number estimation and 100 million per sample for analysis using Hiseq2500.

### Statistical analysis of scRNA-Seq

Quality assessment of scRNA-Seq data and primary analyses were conducted using the Seurat package for R (Butler et al. 2018). Only cells that expressed more than 200 genes were used for further analysis to remove the effects of low-quality cells. Using the FindMarkers function build in Seurat, differentially expressed genes were identified with the threshold for an adjusted P-value at 0.1 in addition to default settings. To identify the post-transcriptionally regulated genes, DEGs were identified in both spliced (Tables S4 and S5) and un-spliced (Tables S6 and S7) forms. For further analysis, only genes showing detectable levels of both spliced and un-spliced forms were selected (Table S8). Pseudo-time analyses were conducted using the monocle package for R (Trapnell et al. 2014; Qiu et al. 2017a, 2017b). RNA velocity analysis was conducted using the RNA velocyto package for python and R (La Manno et al. 2018) with default conditions. To analyze un-spliced RNA expression changes, data for un-spliced transcripts were combined with Seurat data.

### Cell cycle analysis of germ cells

Cell cycle estimation using scRNA-Seq data was conducted using the method previously reported by Macosko *et al*. and Kashima *et al*. (Macosko et al. 2015; Kashima et al. 2018). Five cell cycle phases (G1/S, S, G2, G2/M and M/G1) reflecting gene sets were obtained from Whitfield *et al*. (Whitfield et al. 2002). Genes assigned to the GO term, “meiosis I cell cycle process” were also obtained. From these obtained genes, we excluded those with a low correlation with all genes in the same group in the analyzed cell population (R<0.2). The average expression of the selected genes in each cell was calculated to define the score for each phase. These scores were scaled to obtain a pattern of phase-specific scores for all germ cells. Then, we compared the pattern of phase-specific scores with 13 potential patterns. We generated 6 simple potential patterns (G1/S, S, G2, G2/M, M/G1 and meiosis), all positive, all negative and 5 intermediate patterns (G1/S-S, S-G2, G2-G2/M, G2/M-M/G1 and M/G1-G1/S) using the 75^th^ and 25^th^ percentiles of scores as positive and negative signals, respectively. Each cell was assigned a phase with the highest correlation score. Cells assigned to G1/S and G1/S-S were finally assigned to G1/S, those in S to S, those in S/G2 to S/G2, those in G2 to G2, those in G2-G2/M and G2/M to G2/M, those in G2/M-M/G1 to M, those in M/G1 to M/G1, those in M/G1-G1/S to G1, those in meiosis to meiosis, and those in all positive, all negative, G2/M-M/G1 and M/G1-G1/S to G0.

### Signal activity analysis

To visualize the activity of RB1, p38 MAPK and RA, downstream genes for each signal were obtained from previous report for RB1 and p38 MAPK (Eguchi et al. 2007; Ewen et al. 2010). As the downstream genes of p38 MAPK, we selected 5 genes which showed male enriched expression in germ cells (Ewen et al. 2010). Scaled expression values were averaged for each gene list to calculate the signal activity.

### Gene enrichment analysis

To characterize the major gene expression changes in mutant cells, we utilized Metascape (Zhou et al. 2019) with default conditions and visualize the results using the GOplot package for R (Walter et al. 2015).

### RNA-immunoprecipitation and reverse-transcription quantitative PCR

For RNA immunoprecipitation followed by reverse-transcription quantitative PCR (RT-qPCR) analysis, NANOS2-DND1 expression was induced by Dox (0.5 μg/ml) addition into the conditionally inducible NANOS2-DND1 NIH-3T3 cell line (ciN2D1-3T3), which was established in our laboratory (Hirano et al., manuscript in preparation) in φ10-cm dishes. After collecting cells by trypsin (0.15 mg/ml) treatment for 3 min at 37°C, cells were washed with PBS and lysed in 1 ml of Buffer A (50 mM Tris-HCL (pH 7.4), 150 mM NaCl, 0.5% NP40, 7.5 mM β-glycerophosphate, 1 mM EDTA, 1 mM DTT and 400 units/ml of Recombinant RNase Inhibitor (TaKaRa)) with cOmplete (Roche) on ice for 10 min and spun at 150,000 rpm for 10 min at 4°C. Twenty microliters of anti-FLAG M2 affinity gel (Sigma) was added to the supernatant and incubated for 3 h at 4°C. After 5 washes with Buffer A, co-precipitated RNAs were purified using the RNeasy Mini Kit (Qiagen). After the synthesis of first-strand cDNAs with 200 U of Superscript IV reverse transcriptase (Invitrogen) and 0.25 μg of oligo dT primer, RT-qPCR analyses were carried out with KAPA SYBR FAST qPCR kits using a Thermal Cycler Dice Real Time System (TaKaRa) according to the manufacturer’s instruction. The expression level of mRNA of interest was normalized by *Gapdh*. Then, the fold enrichment of each mRNA in the IP compared with the input was calculated.

### Functional assessment of 3’UTR using the dual reporter system

The 3’UTR of each gene was cloned downstream of the *Emgfp* sequence in the dual reporter vector and the transgenes were introduced into ciN2D1-3T3 cells. One day after transfection of 1 μg of vector using Lipofectamine 2000 (Invitrogen), cells were treated with Dox (0.5 μg/ml) or DMSO for 1 day. Then, cells were collected using RNAiso Plus (TaKaRa) and RNAs were purified. After the synthesis of first-strand cDNAs with 200 U of Superscpript IV reverse transcriptase (Invitrogen) and 0.5 μg of oligo dT primer, RT-qPCR analyses were performed according to manufacturer’s instruction. The expression level of *Emgfp* was normalized by that of *mCherry* and the fold change of each 3’UTR to *Actb* 3’UTR was calculated.

### Materials and Data availability

All unique/stable reagents generated in this study are available from the Lead Contact with a completed Materials and Transfer Agreement.

The scRNA-Seq data generated during this study are included in this article (and its Supplementary Information files). Sequence data will be deposited in the DDBJ (https://www.ddbj.nig.ac.jp/index.html) after the manuscript is accepted for publication.

### Ethical approval

The study reported involves mice. All mice were handled and propagated in accordance with the guidelines of the National Institute of Genetics (NIG), and all experimental procedures were approved by the Committee for Animal Care and Use of NIG.

## Acknowledgements

We thank Drs S. Yamanaka for the anti-DNMT3L antibody, Y. Nishimune for the anti-TRA98 antibody, Atsushi Suzuki for the *Oct4dPE-CreER^t2^*mouse line, Mr. M. Muraoka for his kind support in establishing the analysis environment for chromium data and Danelle Wright for editing this manuscript. We also thank for Dr. A. Toyoda for his kind support to prepare the chromium library and sequencing.

## Author contributions

All experiments except for cell spread staining analysis (conducted by HK) were conducted by RS. The ciN2D1-3T3 cell line was established by TH. The manuscript was written by RS and YS.

