## Supplemental Figure 1 for "NANOS2 suppresses the cell cycle by repressing mTORC1 activators in embryonic male germ cells"

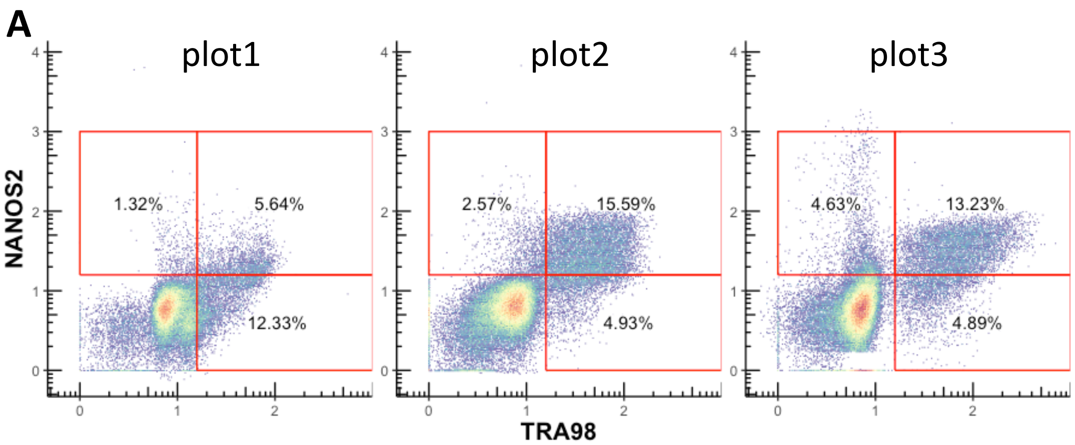


| % | plot1 | plot2 | plo3 | Average ± sd |
|---|---|---|---|---|
| Germ cell | 19.29 | 23.10 | 22.74 | 21.71 ± 2.1 |

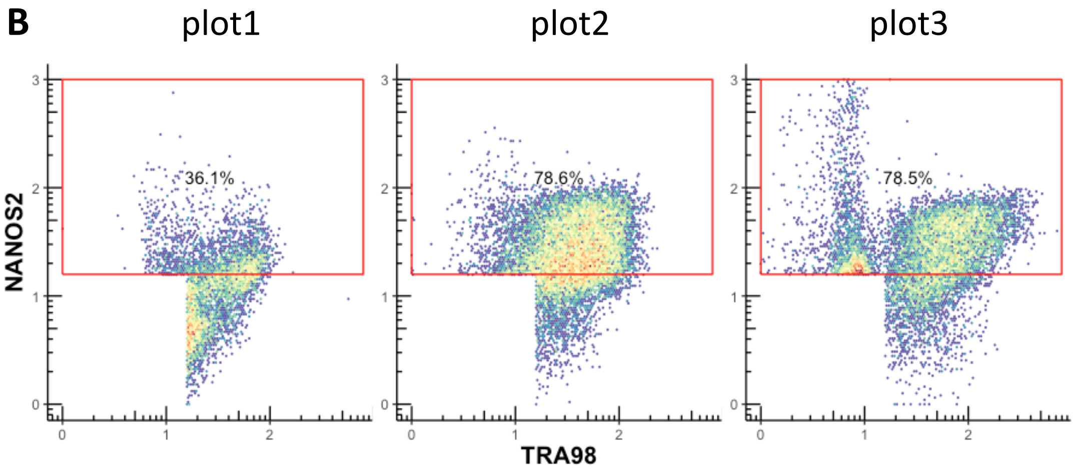


**Figure S1.** FACS analysis for E13.5 testes. Flow cytometric analyses of germ cells obtained from E13.5 testes. Cells were stained with anti-NANOS2 and anti-TRA98 antibodies. The proportion of germ cells was estimated by TRA98-positive and/or NANOS2-positive cells. Each red rectangle indicates each gene-positive cell population estimated from the density plot. (A) contains all cell populations and (B) contains only the germ cell population. Percentages indicate the proportion of selected cells in each plot.

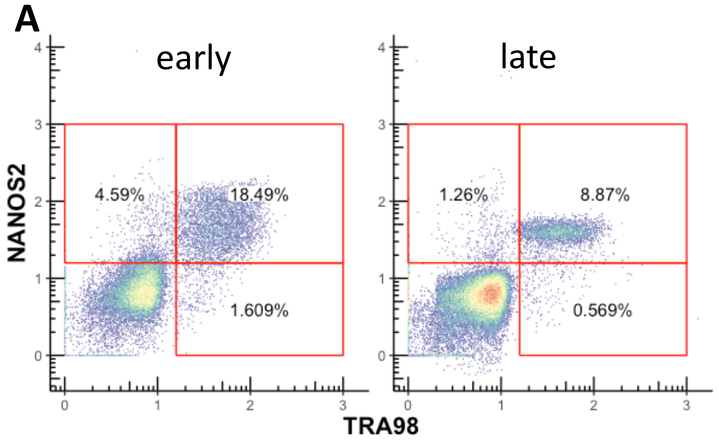


| % | early | late | Average ± sd |
|---|---|---|---|
| Germ cell | 24.7 | 10.7 | 17.7 ± 9.89 |

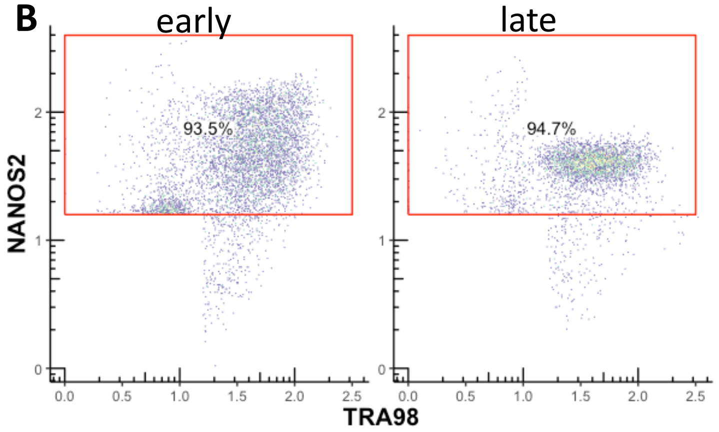


**Figure S2.** FACS analysis for E14.5 testes. Flow cytometric analyses of germ cells obtained from E14.5 testes. The stages of samples were roughly defined based on their embryonic morphology. Cells were stained with anti-NANOS2 and anti-TRA98 antibodies. The proportion of germ cells was estimated by TRA98-positive and/or NANOS2-positive cells. Each red rectangle indicates each gene-positive cell population estimated from the density plot. (A) contains all cell populations and (B) contains only the germ cell population. Percentages indicate the proportion of selected cells in each plot.

A

| MCH male | | E11.5 | E12.5 | E13.5 | E14.5 | E15.5 |
|---|---|---|---|---|---|---|
| Used gonads | | 14 | 6 | 14 | 16 | 10 |
| Cell No. | Total | 2,691 | 6,898 | 2,287 | 606 | 2,588 |
| | After QC | 2,311 (85.9%) | 4,488 (65.1%) | 1,369 (59.9%) | 451 (74.4%) | 1,758 (67.9%) |
| Germ No. | | 102 (4.4%) | 791 (17.6%) | 120 (8.8%) | 54 (12.0%) | 96 (5.5%) |

B

| MCH female | | E11.5 | E12.5 | E13.5 | E14.5 | E15.5 |
|---|---|---|---|---|---|---|
| Used gonads | | 16 | 10 | 18 | 18 | 26 |
| Cell No. | Total | 2,448 | 3,759 | 3,060 | 13,951 | 1,869 |
| | After QC | 2,126 (86.8%) | 2,274 (60.5%) | 1,443 (47.2%) | No data (%) | 1,141 (61.0%) |
| Germ No. | | 76 (3.6%) | 559 (24.6%) | 654 (45.3%) | No data (%) | 220 (19.3%) |

C

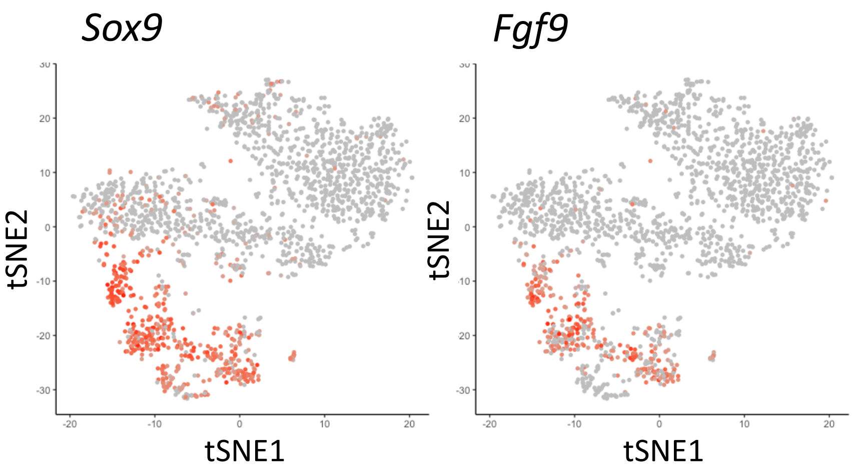


**Figure S3.** Basic information of scRNA-Seq libraries. (A-B) The number of used gonads, sequenced cell number and germ cell number are shown. As an excess number of cells was loaded for E14.5 female, we did not read the sequence deeply (See Methods for detail). (C) Expression levels of male supporting marker genes, *Sox9* and *Fgf9,* are shown in the tSNE plot, which contains all gonadal cells analyzed.

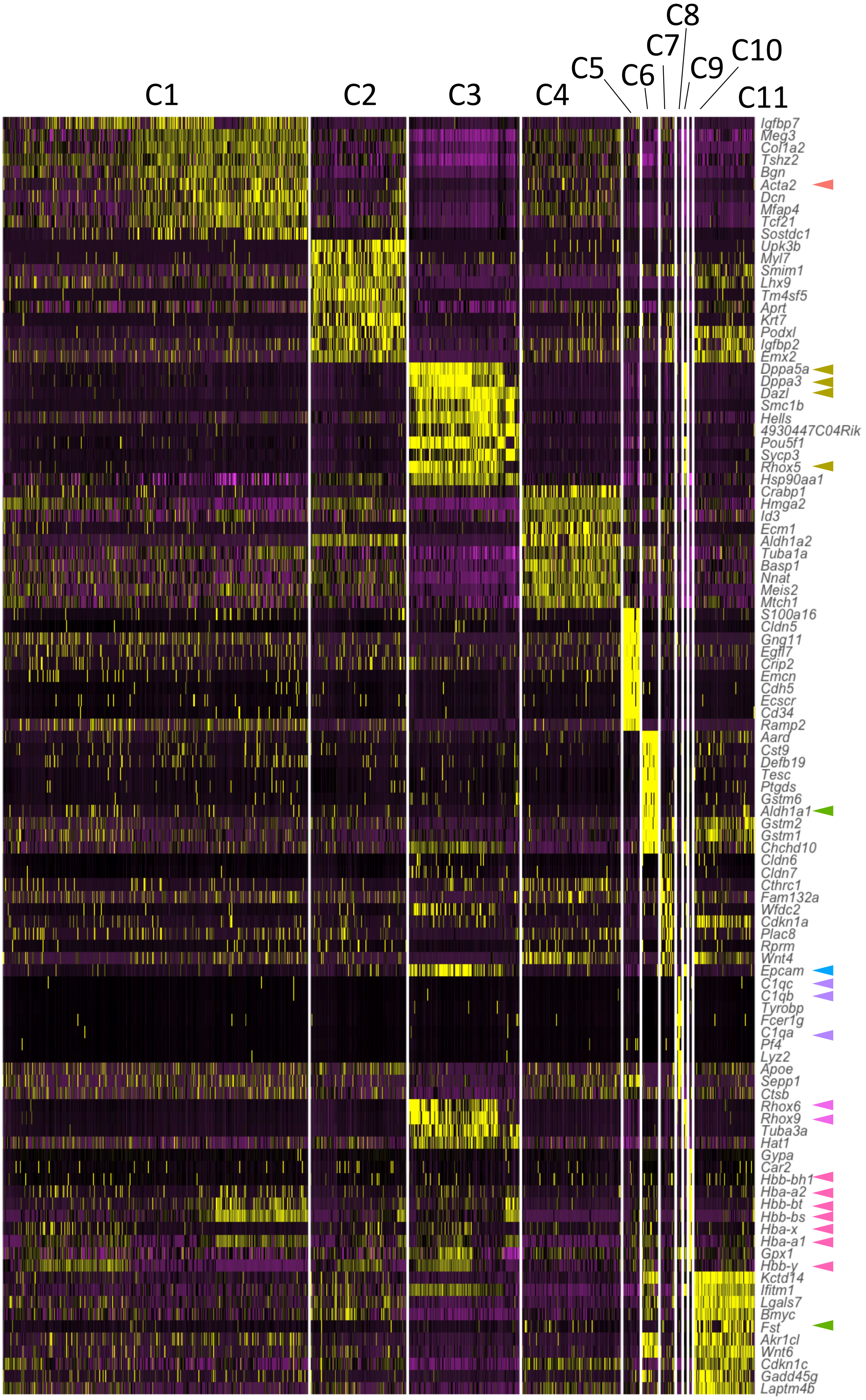
C1
C2
C3
C4
C5
C6
C7
C8
C9
C10
C11
Igfbp7
Meg3
Col1a2
Tshz2
Bgn
Acta2
Dcn
Mfap4
Tcf21
Sostdc1
Upk3b
Myl7
Smim1
Lhx9
Tm4sf5
Aprt
Krt7
Podxl
Igfbp2
Emx2
Dppa5a
Dppa3
Dazl
Smc1b
Hells
4930447C04Rik
Pou5f1
Sycp3
Rhox5
Hsp90aa1
Crabp1
Hmga2
Id3
Ecm1
Aldh1a2
Tuba1a
Basp1
Nnat
Meis2
Mtch1
S100a16
Cldn5
Gng11
Egfl7
Crip2
Emcn
Cdh5
Ecscr
Cd34
Ramp2
Aard
Cst9
Defb19
Tesc
Ptgds
Gstm6
Aldh1a1
Gstm2
Gstm1
Chchd10
Cldn6
Cldn7
Cthrc1
Fam132a
Wfdc2
Cdkn1a
Plac8
Rprm
Wnt4
Epcam
C1qc
C1qb
Tyrobp
Fcer1g
C1qa
Pf4
Lyz2
Apoe
Sepp1
Ctsb
Rhox6
Rhox9
Tuba3a
Hat1
Gypa
Car2
Hbb-bh1
Hba-a2
Hbb-bt
Hbb-bs
Hba-x
Hba-a1
Gpx1
Hbb-y
Kctd14
Ifitm1
Lgals7
Bmyc
Fst
Akr1cl
Wnt6
Cdkn1c
Gadd45g
Laptm4b

**FigureS4.** Heat plot for the top 10 representative genes for each cluster. The top 10 representative genes of each cluster are shown. Yellow indicates strong expression and purple indicates weak expression. Cluster numbers are indicated on the plot. Arrowheads indicate well-known genes that represent specific cell types in gonads. Colors of arrowheads coincide with cell colors in Fig. 2A.

## E13.5

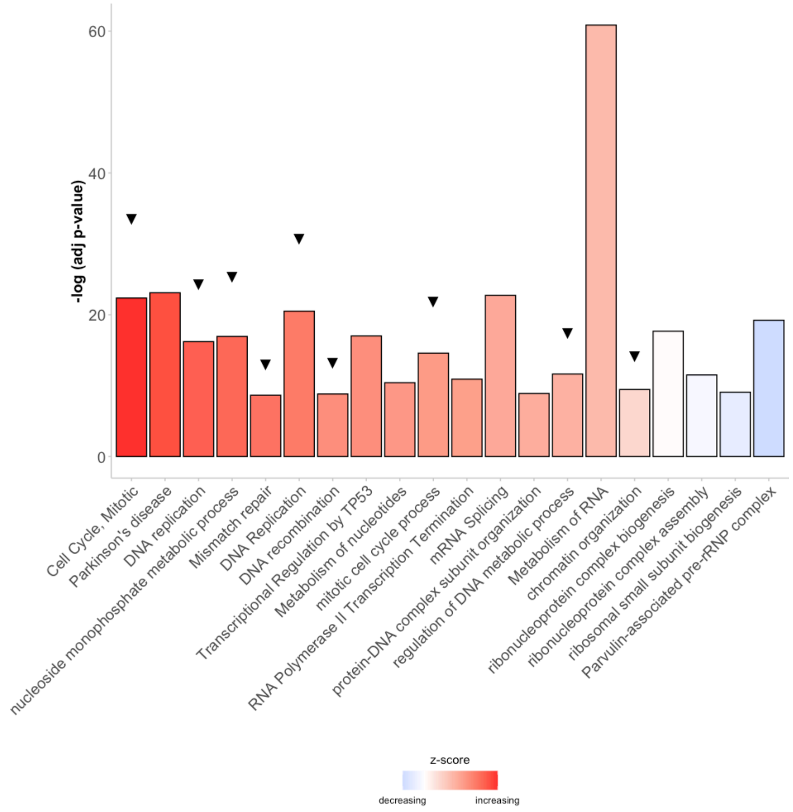
-log (adj p-value)
0
20
40
60
Cell Cycle, Mitotic
Parkinson's disease
DNA replication
nucleoside monophosphate metabolic process
Mismatch repair
DNA Replication
DNA recombination
Transcriptional Regulation by TP53
Metabolism of nucleotides
mitotic cell cycle process
RNA Polymerase II Transcription Termination
mRNA Splicing
protein-DNA complex subunit organization
regulation of DNA metabolic process
Metabolism of RNA
chromatin organization
ribonucleoprotein complex biogenesis
ribonucleoprotein complex assembly
ribosomal small subunit biogenesis
Parvulin-associated pre-rRNP complex
z-score
decreasing
increasing

## E14.5

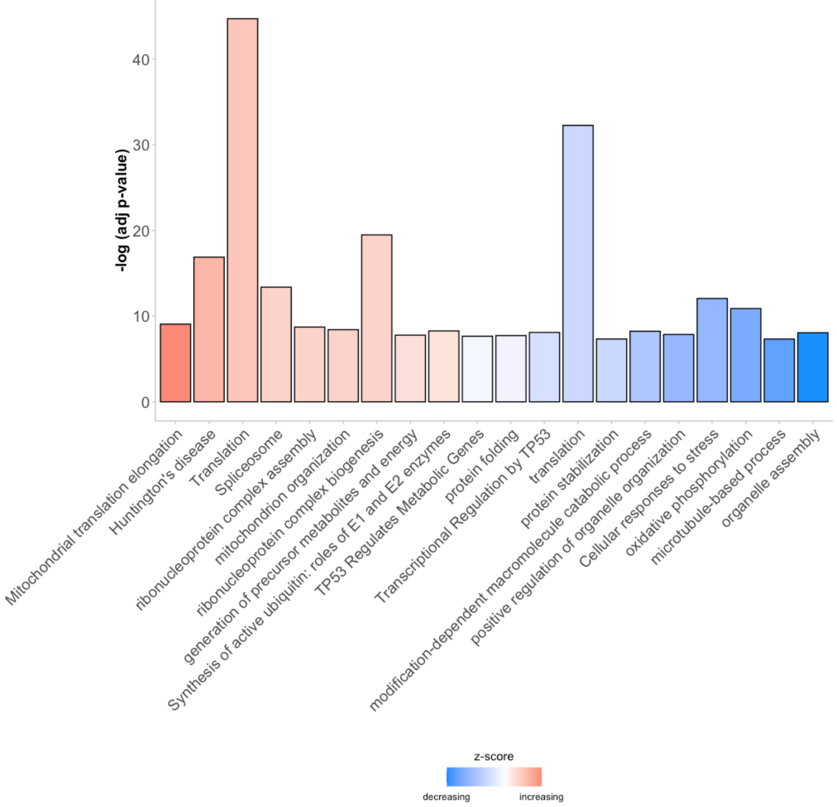
-log (adj p-value)
0
10
20
30
40
Mitochondrial translation elongation
Huntington's disease
Translation
Spliceosome
ribonucleoprotein complex assembly
mitochondrion organization
ribonucleoprotein complex biogenesis
generation of precursor metabolites and energy
Synthesis of active ubiquitin: roles of E1 and E2 enzymes
TP53 Regulates Metabolic Genes
protein folding
Transcriptional Regulation by TP53
translation
protein stabilization
modification-dependent macromolecule catabolic process
positive regulation of organelle organization
Cellular responses to stress
oxidative phosphorylation
microtubule-based process
organelle assembly
z-score
decreasing
increasing

**FigureS5.** Gene enrichment analysis for DEGs in NANOS2-null cells. The result of gene enrichment analysis using Metascape. Heights of each bar indicate the log-transformed adjusted p-value. Colors indicate the z-score, which was calculated as follows: $zscore = \frac{(up-down)}{\sqrt{count}}$. Up, down and count indicate the number of up- or down-regulated and assigned genes in each term from the DEG list, respectively. Black arrowheads indicate cell cycle-related terms.

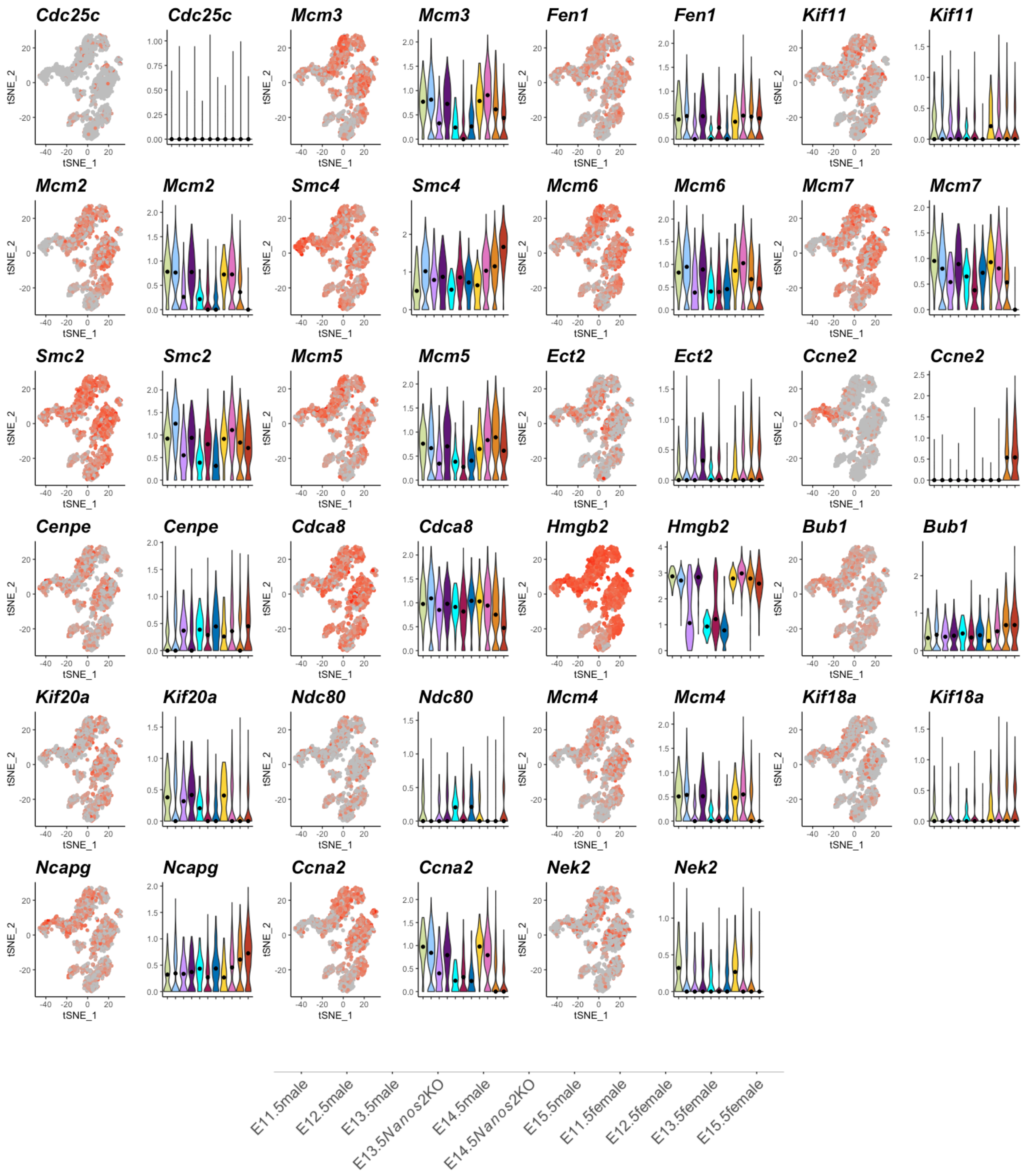


**Figure S6.** Expression patterns of the target genes of E2Fs, which is suppressed by RB1. Expression levels of E2Fs-target genes are visualized on a tSNE-plot (left) and violin plot (right). Black dots indicate the median.

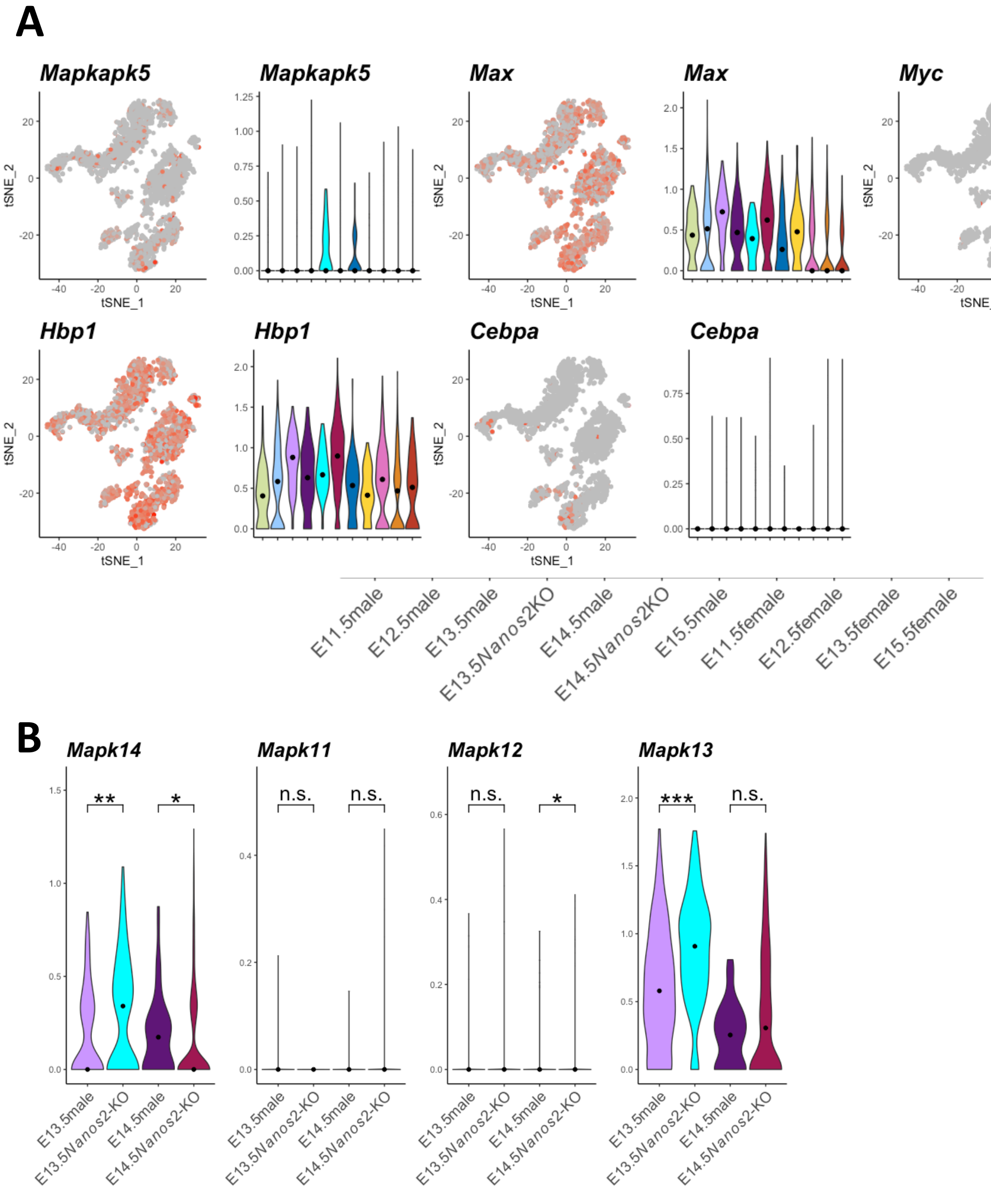


**Figure S7.** Expression patterns of the downstream genes of p38 MAPK. (A) Expression values of p38 MAPK downstream genes are visualized on a tSNE plot (left) and violin plot (right). Black dots indicate the median. (B) Expression values of p38 MAPK genes are visualized on a tSNE plot. Black dots indicate the median. ***P<0.001, **P<0.01, *P<0.05 and n.s.=not significant.

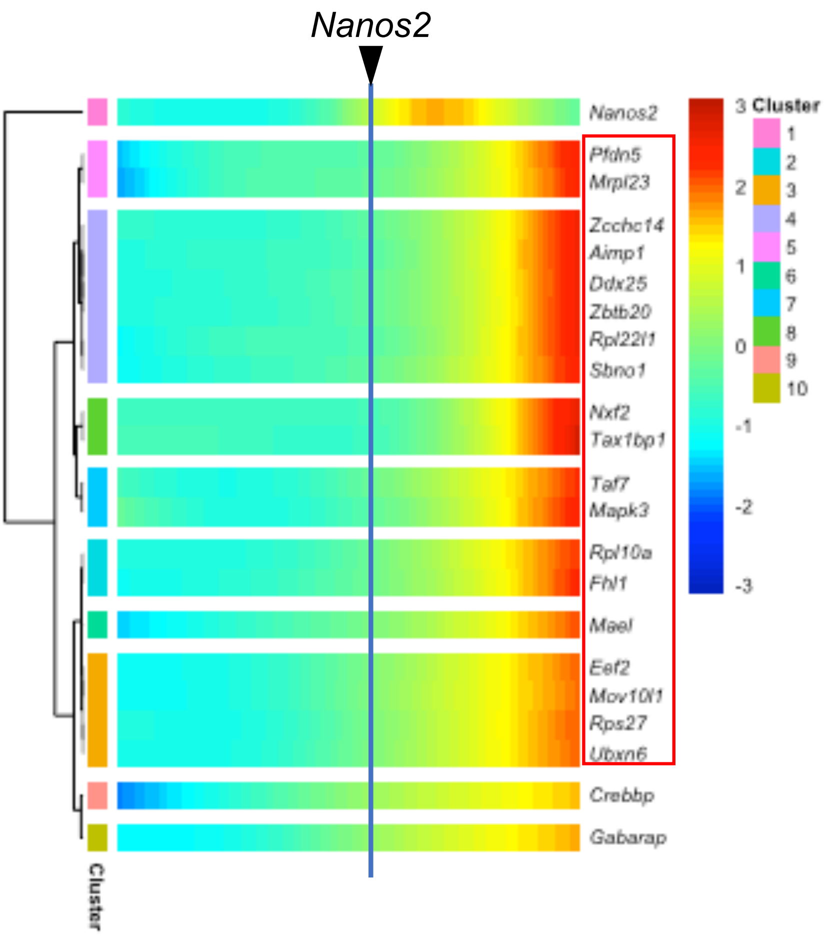


**Figure S8.** Pseudo-time heat plot for upregulated genes. The gene expression patterns of the 21 obtained gene candidates activated by NANOS2 along the developmental pseudo-time of male germ cells. The point when *Nanos2* expression becomes strong is indicated as a vertical line. Genes surrounded by the red rectangle belong to gene groups that were upregulated just after *Nanos2* expression.

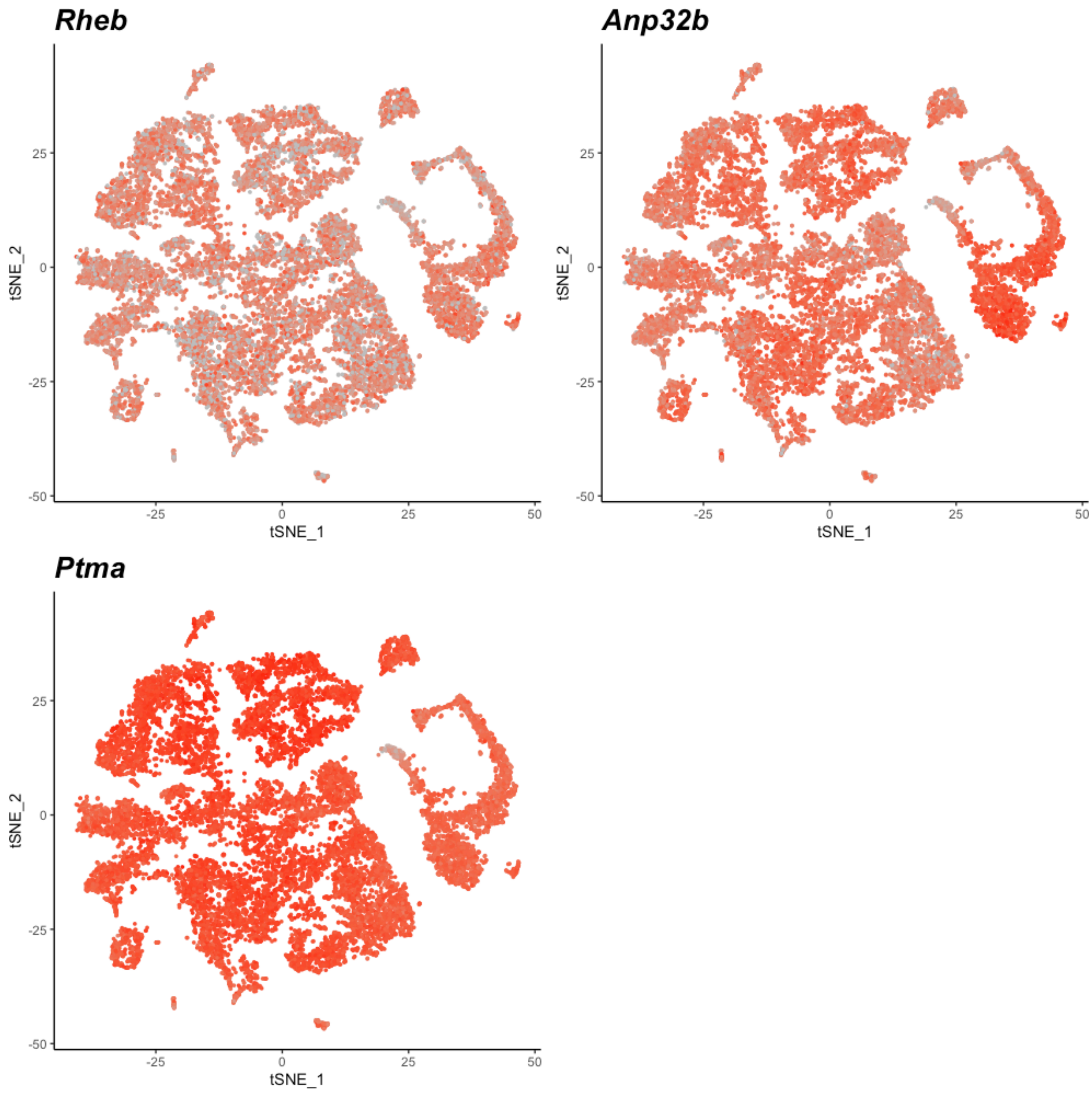


**Figure S9.** Gene expression of obtained gene candidates in developing gonads. Expression values of *Rheb*, *Anp32b* and *Ptma* are shown on the tSNE-plot for all gonadal cells from E11.5-E15.5 embryos. Cells with high expression are colored darker red.

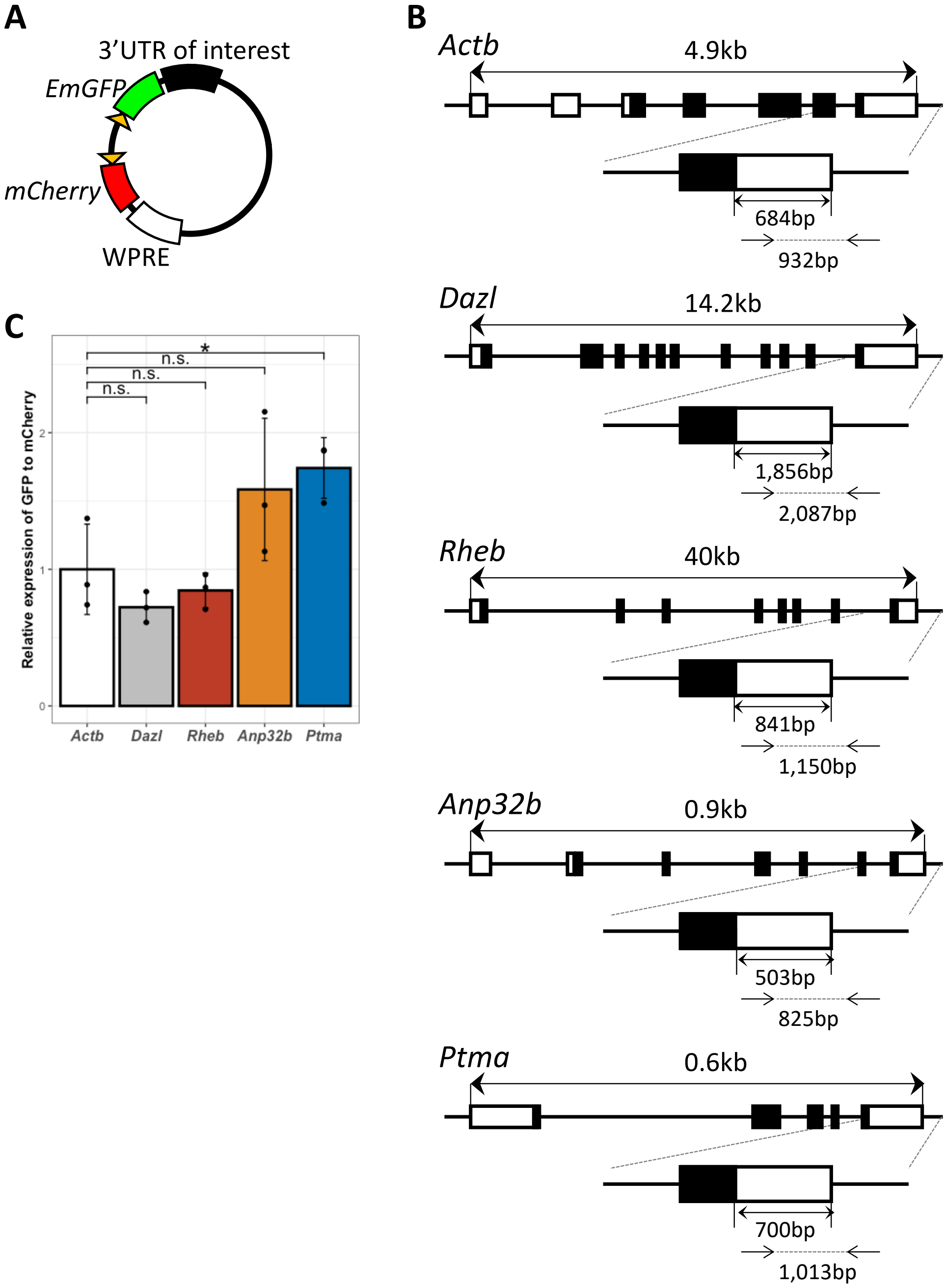


**Figure S10.** Dual-reporter assay for identification of the NANOS2-sensitive 3'UTR. (A) A vector map of the dual fluorescence reporter vector. The internal normalization control *mCherry* and *Emgfp* with the 3'UTR of interest are located in a head-to-head manner and driven by the EF1-α promoter. (B) Gene structures of *Actb*, *Dazl* and 3 gene candidates are shown. The length of the 3'UTR of each gene and inserted portion in the vector are indicated below the gene structure. (C) Relative expression of *Emgfp* in DMSO-treated cells is shown. The *Emgfp* value was normalized by the *mCherry* value, and then relative values to *Actb* 3'UTR were calculated. *P<0.05.
